# Spiking resonances in models with the same slow resonant and fast amplifying currents but different subthreshold dynamic properties

**DOI:** 10.1101/128611

**Authors:** Horacio G. Rotstein

## Abstract

The generation of spiking resonances in neurons (preferred spiking responses to oscillatory inputs) requires the interplay of the intrinsic ionic currents that operate at the subthreshold voltage regime and the spiking mechanism. Combinations of the same types of ionic currents in different parameter regimes may give rise to different types of nonlinearities in the voltage equation (e.g., parabolic- and cubic-like), generating subthreshold oscillations patterns with different properties. We investigate the spiking resonant properties of conductance-based models that are biophysically equivalent at the subthreshold level (same ionic currents), but functionally different (parabolic- and cubic-like). As a case study we consider a model having a persistent sodium current and a hyperpolarization-activated (h-) current. We unfold the concept of spiking resonance into evoked and output spiking resonance. The former focuses on the input frequencies that are able to generate spikes, while the latter focuses on the output spiking frequencies regardless of the input frequency that generated these spikes. A cell can exhibit one or both types of resonance. We also measure spiking phasonance, which is an extension of subthreshold phasonance to the spiking regime. The subthreshold resonant properties of both types of models are communicated to the spiking regime for low enough input amplitudes as the voltage response for the subthreshold resonant frequency band raises above threshold. For higher input amplitudes evoked spiking resonance is no longer present, but output spiking resonance is present primarily in the parabolic-like model, while the cubic-like model shows a better 1:1 entrainment. We use dynamical systems tools to explain the underlying mechanisms and the mechanistic differences between the resonance types. Our results show that the effective time scales that operate at the subthreshold regime to generate intrinsic subthreshold oscillations, mixed-mode oscillations and subthreshold resonance do not necessarily determine the existence of a preferred spiking response to oscillatory inputs in the same frequency band. The results discussed in this paper highlight both the complexity of the suprathreshold responses to oscillatory inputs in neurons having resonant and amplifying currents with different time scales and the fact that the identity of the participating ionic currents is not enough to predict the resulting patterns, but additional dynamic information, captured by the geometric properties of the phase-space diagram, is needed.

## 1 Introduction

Several neuron types have been shown to exhibit preferred frequency responses to oscillatory inputs (resonances) [1–50], which have been implicated in the generation of network oscillations in the same frequency bands [12, 37, 51, 52] (but see [53]). Most studies using single neurons have focused on subthreshold resonance [1–42] and much less attention has been paid to the suprathreshold preferred frequency responses to oscillatory inputs (suprathreshold spiking or firing rate resonance) and the link between the sub- and suprathreshold resonances [2, 22, 38, 43–50, 54]. The mechanisms responsible for the generation of suprathreshold resonance and the circumstances under which the presence of subthreshold resonance is a good predictor of suprathreshold resonance are not well understood. Several studies have suggested that firing resonance emerges from sub-threshold resonance properties [6, 7, 9], but others have not found such a clear correlation [8, 55–57]. This is to be expected, at least in some cases, since suprathreshold resonance depends on the spiking mechanisms in addition to the neuronal intrinsic properties (ionic currents). An important conceptual issue is that, unlike subthreshold resonance, there is more than one notion of the preferred spiking response to oscillatory inputs as we explain below.

The goal of the this paper is to address these issues in the context of nonlinear conductance-based models that are biophysically equivalent at the subthreshold level (same ionic currents), but functionally different in the sense that they have qualitatively different subthreshold oscillatory properties due to differences in parameter values [58]. This allows us to examine various plausible realistic scenarios in which the same participating ionic currents interact both among themselves and with the spiking mechanisms to produce the different types of preferred spiking responses to oscillatory inputs. In addition, it lets us explain the underlying biophysical and dynamic mechanisms in a broader context that goes beyond the interaction between ionic currents. On a more general level, this approach allows to examine the notion of the communication of the resonant properties from the subthreshold to the suprathreshold regime [2, 44, 53].

The subthreshold preferred frequency responses to oscillatory inputs have been characterized by the impedance amplitude (or simply impedance) and phase-shift (or simply phase) profiles (curves of the impedance and phase as a function of the input frequency) [1, 2, 59]. A neuron exhibits subthreshold resonance if the impedance profile peaks at a nonzero input (resonant) frequency (*f_res_*) and subthreshold phasonance if the phase profile vanishes at a nonzero input (phasonant) frequency (*f_phas_*) (the input and output are synchronized in phase). Resonance has being observed in both current and voltage clamp experiments and in models in various neuron types including hippocampal pyramidal cells and interneurons, neocortical neurons, entorhinal stellate cells, thalamic neurons, inferior olive neurons, striatal neurons and pyloric neurons of the crab stomatogastric ganglion neurons [1–42]. Experimental and modeling studies have shown the presence of more complex subthreshold impedance profiles exhibiting additional extrema, referred to as antiresonance (impedance profile minimum) and antiphasonance (zero-phase response with negative slope) [2, 17, 36, 60].

Subthreshold resonance in neurons requires the interplay of positive and negative feedback effects that favor and oppose changes in voltage respectively. These are typically provided by the so-called resonant and amplifying gating variables associated to the corresponding ionic currents. Passive neurons are low-pass filters (monotonically decreasing impedance profile and monotonically increasing phase profile with no zero-crossing). The presence of restorative ionic currents such as *I_h_* (hyperpolarization-activated mixed-cation) and *I_M_* (M-type slow-potassium) endows neurons with the ability to exhibit resonance, while the presence of regenerative currents such as *I_Nap_* (persistent sodium) and *I_Kir_* (inward-rectifying potassium) amplify the neurons’ response to oscillatory input in addition to other causing changes in the impedance and phase profiles [1, 2, 59, 61]. Negative feedback effects must be slower than the fastest regenerative process for resonance to occur.

When measuring subthreshold resonance in single neurons one typically (and often implicitly) assumes that both the input and output frequencies coincide and the response amplitude is uniform across cycles for a given input frequency. This lack of harmonics allows the impedance profile to capture the amplitude of the voltage response and *f_res_* to be a predictor of what input frequencies will produce spikes for low enough suprathreshold input amplitudes. In this sense, it is often said that subthreshold resonance is communicated to the suprathreshold regime [2] and has implications for the the generation of neuronal oscillations [1]. However, for larger input amplitudes the input frequency band that is able to produce spikes expands and may do so away from the boundaries of the underlying subthreshold resonant frequency band (e.g., theta: 4 − 10 Hz). The neuron may even become a spiking low-pass filter where all input frequencies below some limit are able to produce spikes. On the other hand, since spiking may skip input cycles for a given frequency, the output frequency may remain bounded within a given frequency band for input frequencies in much larger ranges.

We use the term resonance as a synonym for frequency preference response to oscillatory inputs and unfold the concept of spiking resonance into evoked and output spiking resonance. The former focuses on the input frequencies that are able to generate spikes [43], while the latter focuses on the output spiking frequencies regardless of the input frequency that generated these spikes. A cell can exhibit one or both types of resonance. A related preferred frequency response is spiking phasonance, which is an extension of subthreshold phasonance to the spiking regime. The circumstances under which these resonances occur and coexist, and their dependence on the intrinsic ionic currents and their interaction with the oscillatory inputs are not well understood.

In previous work we investigated the mechanisms of generation of subthreshold oscillations (STOs) in conductance-based models whose subthreshold dynamics are described by the same combinations of ionic currents (*I_h_* + *I_Nap_* and *I_Ks_* + *I_Nap_*, both including a leak current), but give rise to different types of nonlinearities (parabolic-like and cubic-like) in different biophysically plausible parameter regimes [58]. These models have been used to investigate the generation of STOs and mixed-mode oscillations (MMOs) in entorhinal cortex stellate cells [61–64]. We showed that while some STO properties are controlled by the specific types of ionic currents involved, and they are different for the *I_Nap_* + *I_h_* and the *I_Nap_* + *I_Ks_* models, other properties are controlled by the geometry of the phase-plane and are shared by models with different ionic currents, but the same type of voltage nullclines (parabolic- or cubic-like). The question arises whether, and if yes how and under what conditions, these similarities and differences are reflected in the spiking response of models with the same ionic currents and qualitatively different phase-space diagrams.

The outline of the paper is as follows. In Section 3.1 we discuss the intrinsic STO and spiking patterns that arise in the parabolic- and cubic-like *I_h_* + *I_Nap_* models (in the absence of any structured oscillatory input) and we examine the similarities and differences in the mechanisms underlying the generation of these patterns between the two types of models. In the presence of noise the intrinsic dynamics of these two models are almost indistinguishable. In Section 3.2 we show that the subthreshold voltage responses of the two models have different gain dependencies for large enough (subthreshold) input amplitudes. As compared to the corresponding linearized models, the impedance amplitude is larger for the parabolic-like model and smaller for the cubic-like model. In Section 3.3 we define and characterize the three types of spiking responses we use to investigate suprathreshold resonance: evoked, output and phase responses. Evoked spiking resonance occurs when spiking is generated only for input frequencies in an intermediate (resonant) frequency band regardless of the output frequency. Output spiking resonance occurs when the output frequency belongs to an intermediate (resonant) frequency band regardless of the input frequency. Spiking phasonance is an extension of subthreshold phasonance. It occurs when the phase-shift (or phase) between the output spike and the input peak vanishes at a non-zero input frequency. In Section 3.4 we examine the similarities and differences in the mechanisms underlying the generation evoked, output and phase resonance between the two models. Both exhibit evoked and output spiking resonance for low enough values of (suprathreshold) input amplitudes. However, evoked spiking resonance vanishes for larger values of the input amplitude. Output resonance persists in the parabolic-like model, but not in the cubic-like model. This critically depends on the ability of the parabolic model to generate response mixed-mode oscillatory patterns, which is almost absent in the cubic-like model. Finally, we discuss our results, extensions for other time constant regimes, limitations and implications for neuronal dynamics in Section 4.

## 2 Methods

### 2.1 Conductance-based *I_h_* + *I_Nap_* models

We use conductance-based models of Hodgkin-Huxley type [65] whose subthreshold dynamics involve the interplay of three ionic currents: passive leak (*I_L_*), hyperpolarization-activated or h-(*I_h_*), and persistent sodium (*I_Nap_*). We refer generically to these models as *I_h_* + *I_Nap_*. These models do not include the spiking currents (transient sodium and delayed-rectifier potassium) and therefore they do not describe the spike dynamics. Spikes are added “manually” after their onset has been detected. The latter either results from the model subthreshold dynamics (parabolic-like model 1) or by a voltage threshold mechanism (cubic-like model 2) as in the standard models of integrate- or resonate-and fire type.

The current-balance equation is given by

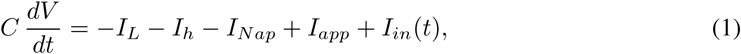
where *V* is the membrane potential (mV), *t* is time (ms), *C* is the membrane capacitance (*μ*F/cm^2^), *I_app_* is the applied bias (DC) current (*μ*A/cm^2^), *I_in_*(*t*) is a time-dependent input current (*μ*A/cm^2^), and the ionic currents are described

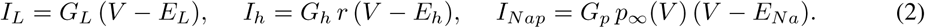

In (2), *r* and *p* are the gating variables, *G_j_* (*j* = *h, p, L*) are the maximal conductances (mS/cm^2^), and *E_j_* (*j* = *h, p, L*) are the reversal potentials (mV). The gating variables *x* (= *r, p*) obey first order differential equations of the form

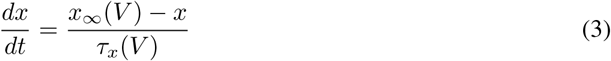
where *x_∞_*(*V*) and *τ_x_*(*V*) are the voltage-dependent activation/inactivation curves and time-scales respectively. The gating variable *p* for *I_Nap_* in (2) is typically very fast, and it is assumed here to be slave to voltage: *p* = *p_∞_*(*V*). The activation and inactivation curves for *I_Nap_* and *I_h_* are given, respectively, by

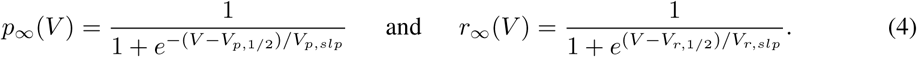

The time constant for *I_h_* is given by *τ_r_* = 80 ms. In the following we will omit the units unless necessary for clarity.

For the sinusoidal inputs with frequency *f_in_* (Hz) we use the following notation

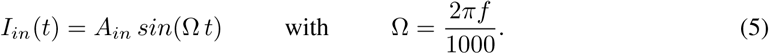

When necessary for clarity, in the graphs we will use the notation *f_in_* for the input frequency.

For some of the simulations of the unforced system we added white noise to the model. Specifically, we added a stochastic term of the form 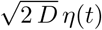 to the right hand side of eq. (1). This term is delta correlated with zero mean; i.e., < *η*(*t*), *η*(*t′*) >= *δ*(*t* – *t′*). *D* > 0 is the standard deviation.

### 2.2 Two *I_h_* + *I_Nap_* models with different geometric/dynamic properties

The two models we use in this paper have the same ionic currents but different parameter values that endow them with qualitatively different geometric/dynamic properties reflected in the shapes of their voltage nullclines in the phase-plane diagram [58]. The *I_h_* + *I_Nap_* model 1 has a parabolic-like *V*-nullcline (Fig. 3-A1) and the *I_h_* + *I_Nap_* model 2 has a cubic-like *V*-nullcline (Fig. 3-B1). Therefore, we will often refer to them as the parabolic and cubic *I_h_* + *I_Nap_* models, the *I_h_* + *I_Nap_* models in the parabolic and cubic regimes, or, simply, the models 1 and 2, respectively. Both are modifications of models originally used for medial entorhinal cortex layer II stellate cells (see below) and are representative of a general class of models having combinations of these currents. The activation/inactivation curves for the two models (blue for model 1 and red for model 2) are presented in Fig. 1.

**Figure 1:**
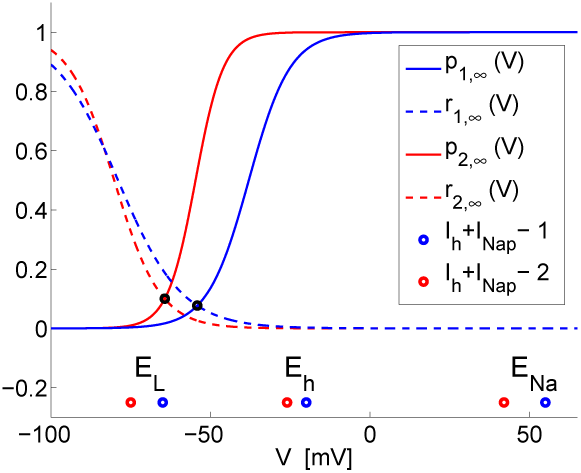
Activation curves for the *I_h_*+*I_Nap_* models 1 and 2. *The subindex j* = 1, 2 *in p_j,∞_(V) and r_j,∞_(V) refers to the models 1 and 2 accordingly. The blue and red dots indicate the values of the reversal potentials E_L_, E_h_ and E_Na_ for models 1 and 2 respectively*.

**Figure 2:**
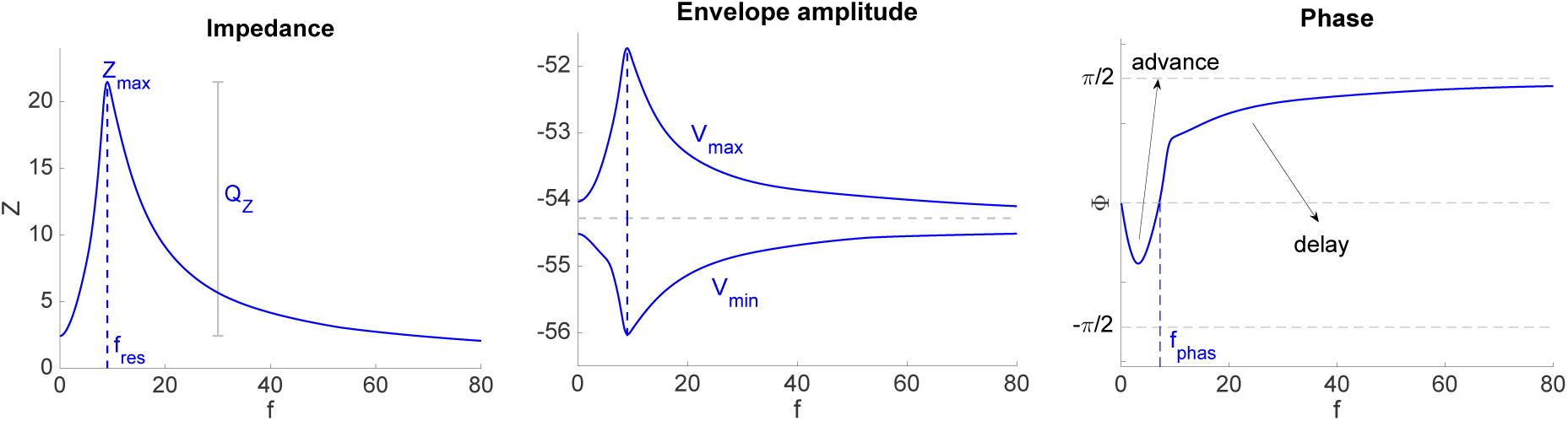
Representative Impedance (*Z*), envelope amplitude and phase (Φ) profiles. *The impedance and phase profiles are the curves of Z and* Φ *vs. the input frequency f. The envelope amplitude are the curves of the maximum (V_max_) and minimum (V_min_) voltage values of the steady state response. Resonance occurs at f* = *f_res_. The resonance amplitude Q_z_* = *Z_max_* − *Z*(0). *Phasonance occurs at f* = *f_phas_. The voltage response is advanced for f* < *f_phas_ and delayed for f* > *f_phas_. The lack of symmetry between the upper and lower branches of the envelope amplitude response reflects the nonlinearities present in the system*.

**Figure 3:**
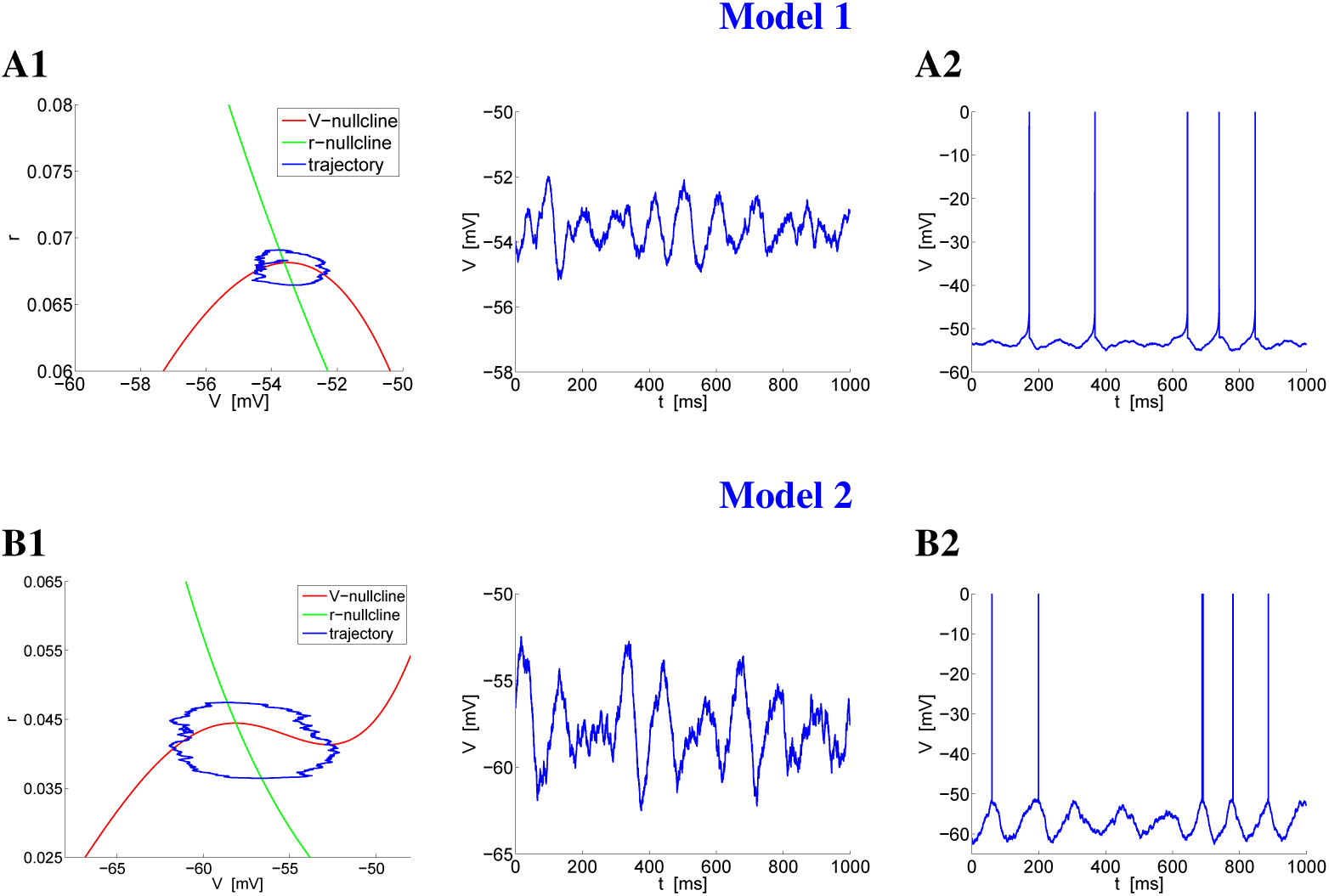
STOs and spiking dynamics for the autonomous *I_h_*+*I_Nap_* models 1 (A; parabolic regime) and 2 (B; cubic regime) for representative parameter values. *In the absence of noise (D* = 0*) both models have a stable fixed-point. For visualization purposes, only one STO cycle is shown in the phase-plane diagrams (A1 and B1, left) and spikes were truncated (A2 and B2)*. **A.** *I_h_*+*I_Nap_* **model 1.** *We used the following parameter values: G_L_* = 0.5, *G_p_* = 0.5, *G_h_* = 1.5, *I_app_* = −2.25 *(A1), *I_app_** = −2.2 *(A2), D* = 0.004, *V_th_* = −50, *V_rst_* = −52, *r_rst_* = 0.065. **B.** *I_h_*+*I_Nap_* **models 2.** *We used the following parameter values: G_L_* = 0.3, *G_p_* = 0.08, *G_h_* = 1.5, *I_app_* = 0.4 *(B1), *I_app_** = 0.55 *(B2), D* = 0.04, *V_th_* = −51, *V_rst_* = −52, *r_rst_* = 0.035.

#### *I_h_* + *I_Nap_* model 1

This model is a modified version [61, 62] of the one model introduced in [66]. The spiking currents were eliminated without affecting the mechanism that governs the onset of spikes [62], and spikes were artificially reintroduced [61, 62], as for the 2D models of quadratic integrate-and-fire type [67, 68] that have a parabolic voltage nullcline and an additional recovery variable that captures the effects of restorative currents. An additional slow h-current was also eliminated as in [61]. The ability of the model to produce subthreshold oscillations (STOs) and spikes is not affected by these modifications (Fig. 3-A), but the model cannot produce mixed-mode oscillations (MMOs, consisting of STOs interspersed with spikes) in the absence of noise (Fig. 3-A3) or a time-dependent input. For intrinsic mixed-mode oscillations to occur 3D subthreshold dynamics are necessary. We use the following baseline parameter values unless indicated otherwise [62, 66, 69]: *V*_*p*,1/2_ = −38, *V*_*p,slp*_ = 6.5, *V*_*r*,1/2_ = −79, *V*_*r,slp*_ = 10, *C* = 1, *E_L_* = −65, *E_Na_* = 55, *E_h_* = −20, *G_L_* = 0.5, *G_p_* = 0.5 and *G_h_* = 1.5.

#### *I_h_* + *I_Nap_* model 2

This model is a modified version of the model introduced in [64]. We have translated the two nullclines to lower voltage values so the subthreshold voltage regime is in a similar range as for model 1. In contrast to the latter, the spiking mechanism is implemented by adding an artificial voltage threshold (*V_th_*) and a reset mechanism (*V_rst_* and *r_rst_*). We use the following baseline parameter values unless indicated otherwise: *V*_*p*,1/2_ = −54.8, *V*_*p*,*slp*_ = 4.4, *V*_*r*,1/2_ = −74.2, *V*_*r*,*slp*_ = 17.2, *C* = 1, *E_L_* = −75, *E_Na_* = 42, *E_h_* = −26, *G_L_* = 0.3, *G_p_* = 0.08 and *G_h_* = 1.5.

### 2.3 Subthreshold impedance and phase profiles in response to oscillatory input currents: resonance and phasonance

The voltage response of a neuron to oscillatory input currents of the form (5) with amplitude *A_in_* can be characterized by the so-called impedance (*Z*) and phase (Φ) profiles (Fig. 2), which are curves of the impedance amplitude (or simply impedance) and phase-shift (or simply phase) as a function of the input frequency (*f*), defined as [61]

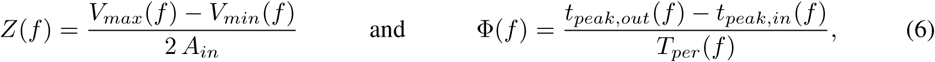
respectively, where *V_max_* (*f*) and *V_min_* (*f*) are the maximum and minimum of the oscillatory voltage response *V_out_*(*f*) respectively, *t_peak,in_* (*f*) and *t_peak,out_* (*f*) are the peak times for the current input and voltage output respectively, and *T_per_* (*f*) is the oscillation period. We note that *Z* (*f*) is often used to refer to the complex impedance (a quantity that includes both amplitude and phase); following other authors we use *Z* (*f*) rather than |*Z*(*f*)| for the impedance amplitude.

Eqs. (6) generalize the impedance and phase profiles for linear systems [2,59] under the assumption that the number of input and output cycles per unit of time coincides and the output voltage waveforms for each each input frequency are identical across cycles (assuming steady state). This is the case for linear (or linearized) and quasi-linear models [59, 61, 70] and the parabolic- and cubic-like models we use in this paper. We note that in other parameter regimes, the above-mentioned assumptions may not be satisfied for cubic-like models.

A neuron exhibits subthreshold *resonance* if *Z*(*f*) peaks at a non-zero (resonant) frequency *f_res_* and subthreshold *phasonance* if the Φ = 0 at a non-zero (phasonant) frequency (*f_phas_*). We expand below

### 2.4 Suprathreshold spike-frequency and spike-phase diagrams in response to oscillator input currents

Here we focus on two measures of the neuronal suprathreshold response to oscillatory input currents: spike-frequency *f_spk_* and spike-phase Φ*_spk_*.

The spike-frequency *f_spk_* was computed as the inverse of the average interspike-intervals (ISI) over 1000 ms (Hz). For the purposes of this paper this measure is appropriate to capture the desired behavior of both types of models and the differences between them. A more detailed analysis (beyond the scope of this paper) that uses noise in addition to oscillatory inputs will require the development of more advanced measures. The measure we use here is closer to the time average rate used in [9, 17] than to the signal gain used in [2] (an extension of the impedance to the suprathreshold regime, defined as the quotient between the instantaneous firing rate of a population of neurons driven by noise as well as a common oscillatory input and the amplitude of this input). The latter requires an oscillatory input with small enough amplitude so that the trial-averaged instantaneous can be captured by a linear suprathreshold response. In contrast, here we are interested in the spiking responses away from the weak input amplitude regime.

The spike-phase (spike-phase-shift) Φ_*spk*_ was computed in a similar manner as the subthreshold phase (phase-shift) in (6) with *t*_*peak*,*out*_ replaced by the spike time and averaged over 1000 ms. Each input cycle was considered to begin and end at the immediate consecutive troughs. Spikes occurring at the peak of the input oscillation cycle were assigned Φ_*spk*_ = 0. Spikes occurring at the immediate prior and posterior troughs were assigned Φ_*spk*_ = ±0.5 and Φ_*spk*_ = −0.5 respectively.

### 2.5 Numerical simulation

The numerical solutions were computed by using the modified Euler method (Runge-Kutta, order 2) [71] with a time step Δ*t* = 0.1 ms in MATLAB (The Mathworks, Natick, MA).

## 3 Results

### 3.1 Intrinsic dynamics: similarities and differences between the parabolic- and cubic-like *I_h_* + *I_Nap_* models

Fig. 3 illustrates the subthreshold (panels A1 and B1) and mixed-mode oscillatory (MMO) dynamics (panels A2 and B2) for the *I_h_* + *I_Nap_* models 1 (panels A) and 2 (panels B) in the theta frequency band in the presence of additive white noise (standard deviation *D*) in the current-balance equation (1). The values of *D* were adjusted in each case in order to illustrate the characteristic patterns.

Geometrically, the most prominent qualitative difference between the *I_h_* + *I_Nap_* models 1 and 2 are captured by the shapes of the corresponding *V*-nullclines [58] given by

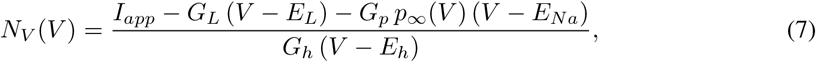
which are parabolic-like for the model 1 (Fig. 3-A1) and cubic-like for the model 2 (Fig. 3-B1). While there are also differences in the r-nullclines *N_r_*(*V*) = *r_∞_*(*V*) between the two models, these are relatively minor.

These differences in the dynamic structure between the two models have consequences not only for the STO dynamics, as shown in [58], but also for the subthreshold and spiking resonant properties as we show below in this paper.

#### 3.1.1 Intrinsic subthreshold oscillations (STOs)

The similarities and differences in the mechanisms of generation of STOs in the parabolic- and cubic-like *I_h_*+*I_Nap_* as well as the mechanisms of generation of STOs and the transition from STOs to spikes in the *I_h_*+*I_Nap_* model 1 were thoroughly analyzed in previous work [58, 62, 63] (see also [61]). The fixed-point in Fig. 3-A1 is a stable focus. In the absence of noise (*D* = 0) model 1 can generate damped STOs (Fig. 5-A2). Persistent STOs (Fig. 3-A1) emerge when white noise is added (*D* > 0). The transition from STOs to spiking as *I_app_* increases occurs through a sub-critical Hopf bifurcation as the *V*-nullcline shifts down and the fixed-point moves to the right relative to the peak of the *V*-nullcline (Fig. 5-A). Alternatively, spiking can be created by noise, as in Fig. 3-A2, when the fixed-point is stable.

**Figure 4:**
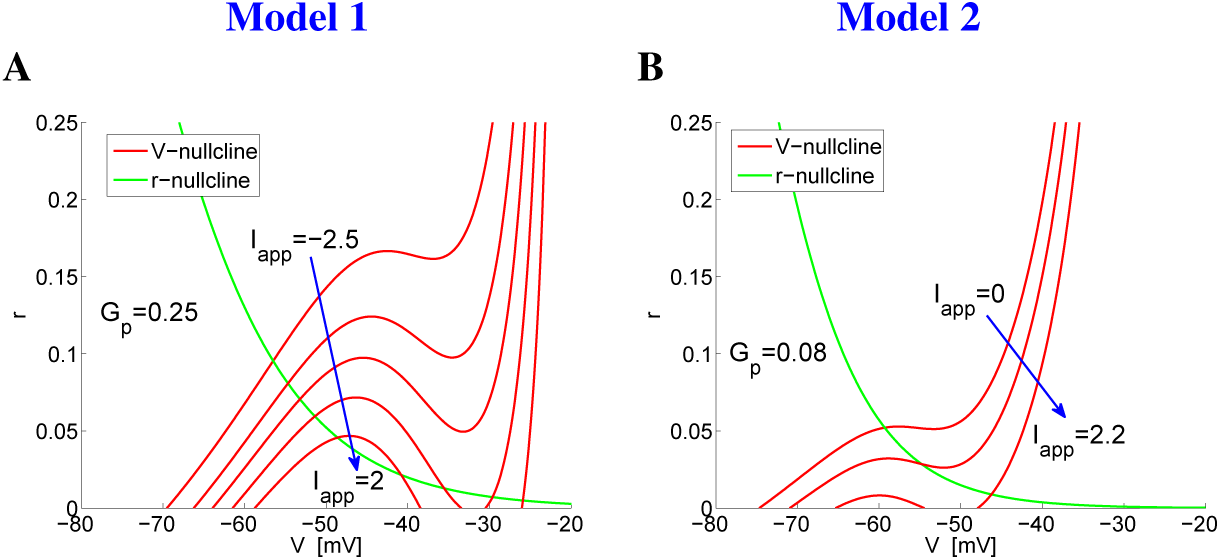
Dependence of the *V*-nullcline on *I_app_* for the autonomous *I_h_*+*I_Nap_* models 1 (A) and 2 (B) for representative parameter values. *The arrow in the phase-plane diagrams indicates the direction of increasing values of *I_app_*. We used the following parameter values: G_L_* = 0.5, *G_h_* = 1.5 *(model 1) and G_L_* = 0.3, *G_h_* = 1.5 *(model 2)*.

**Figure 5:**
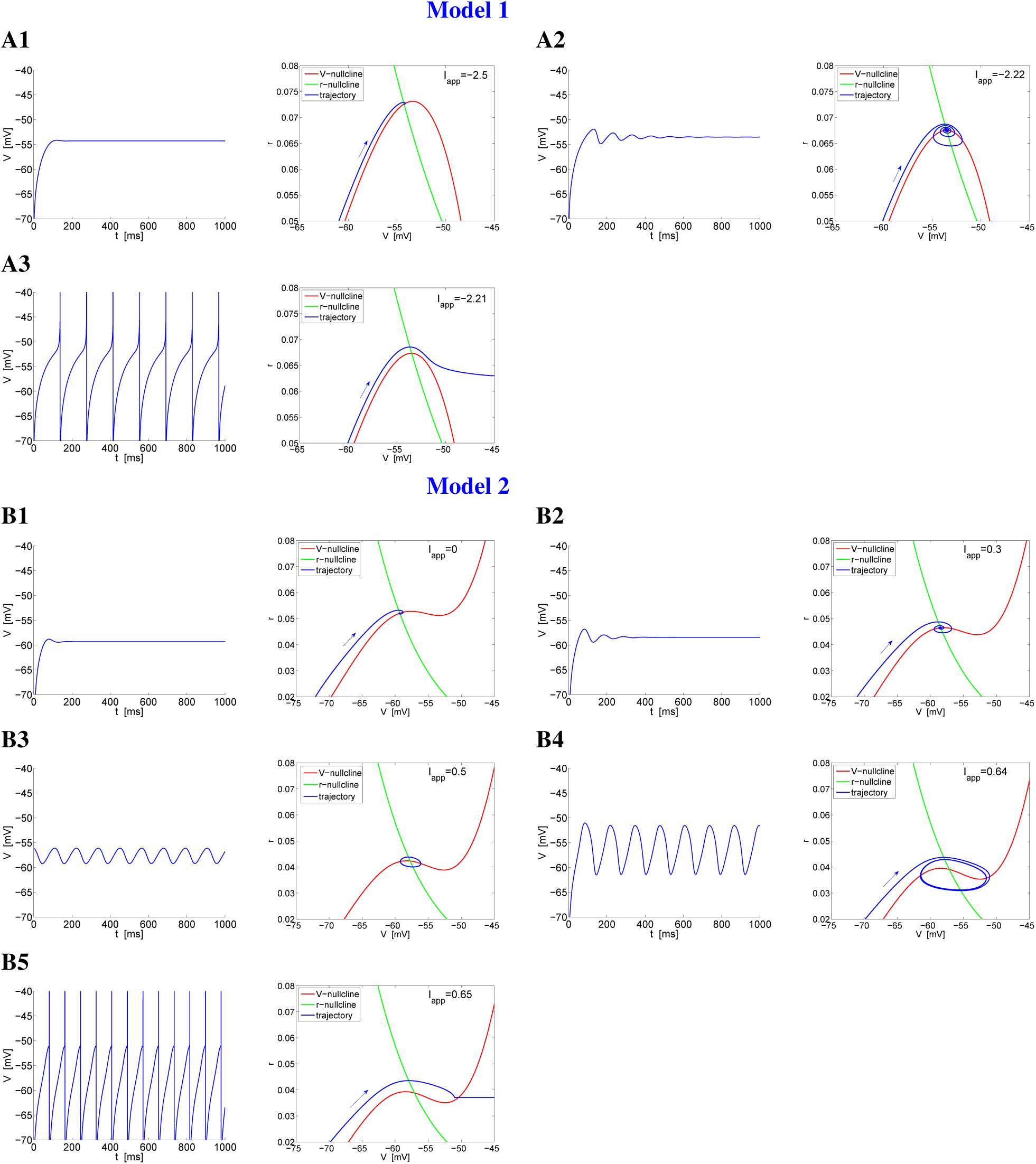
Spiking mechanisms in the autonomous *I_h_*+*I_Nap_* models 1 (A) and 2 (B) for representative parameter values: effects of changes in *I_app_*. **A.** *I_h_*+*I_Nap_* **model 1.** *We used the following parameter values: G_L_* = 0.5, *G_p_* = 0.5, *G_h_* = 1.5, *D* = 0, *V_th_* =?45, *V_rst_* = −75, *r_rst_* = 0.0 **B.** *I_h_*+*I_Nap_* **models 2.** *We used the following parameter values: G_L_* = 0.3, *G_p_* = 0.08, *G_h_* = 1.5, *D* = 0, *V_th_* = −51, *V_rst_* = −75, *r_rst_* = 0

Because the system is fast-slow, for small enough values of *D* the trajectories can move around the knee of the *V*-nullcline (see Section 3.1.2, below). This includes their canard-related ability to move along the unstable branch (right) of the parabolic-like V-nullcline for a significant amount of time before either crossing it and generating a STO (Fg. 5-A2) or moving to the right along a fast fiber into the spiking regime (Fig. 5-A3).

In contrast to model 1, model 2 can generate persistent STOs (Figs. 5-B3 and -B4) in the absence of noise (*D* = 0) in addition to damped oscillations (Figs. 5-B1 and -B2). Although the time scale separation between *V* and *r* is the same (*τ_r_* = 80) for both models, it affects the dynamics of the model 2 in the vicinity of the knee of the parabolic-like nullcline in a different way than for model 1. Specifically, the stable small amplitude oscillations (Fig. 5-B3) generated in the supercritical Hopf bifurcation gradually grow in size until they reach the left branch of the cubic-like *V*-nullcline (Fig. 5-B4). Because of the small distance between nullcline’s local extrema relative to the *I_h_* time constant, these oscillations are not of relaxation type. Finally, although the model includes a sodium current, the subthreshold dynamics does not posses a mechanism for the onset of spikes. As for other cubic-like systems, the same vector field that gives rise to the STOs prevents the trajectories from escaping to the subthreshold voltage regime. Spiking is created by a threshold mechanism as explained earlier in the paper.

#### 3.1.2 Intrinsic spiking mechanisms

Here we examine the mechanisms of transition between STOs and spikes in the autonomous *I_h_*+*l_Nap_* models 1 and 2 as *I_app_* changes. The differences between the two models result from the qualitatively different effects that changes in the excitability levels (*I_app_*) have on the shapes of their respective *V*-nullclines (see [61] for a detailed discussion).

For model 1, as *I_app_* increases the *V*-nullcline shifts down and the fixed-point moves to the right (larger values of *V*). The shape of the *V*-nullcline remains almost unchanged. For low enough values of *I_app_* the fixed-point is a stable node (not shown). The fixed-point transitions to a stable focus as it gets closer to the knee of the *V*-nullcline (Figs. 5-A1 and -A2). The amplitude of the STOs increases with increasing values of *I_app_*. The transition from STOs to spikes occurs through a subcritical Hopf bifurcation (type II excitability; see [72]) and involves the subcritical canard phenomenon [62, 63, 73].

The limit cycle trajectory is able to move along the unstable (right) branch of the *V*-nullcline before crossing it and turning left towards the stable (left) branch, which is a signature of the canard phenomenon. The fixed-point in Fig. 5-A3 is still a stable focus, separated from the trajectory by an unstable small amplitude limit cycle (not shown) [62]. The spiking trajectory first moves around this small amplitude unstable limit cycle and then moves to the right (in the direction of increasing values of *V*) along a fast fiber towards the spiking regime. The voltage threshold indicates that a spike has occurred, but is not part of the spike generating mechanism. The canard explosion of the unstable limit cycle as *I_app_* decreases from the values in Figs. 5-A3 to Figs. 5-A2 allows trajectories initially far away (e.g., reset values for *V* and *r*) to reach the stable fixed-point. As *I_app_* continues to increase, the *V*-nullcline continues to shift down, the fixed-point loses stability, and eventually disappears in a saddle-node bifurcation on the right branch (not shown).

For model 2, increasing values of *I_app_* cause a deformation in the shape of the *V*-nullcline, an overall shifting down of the *V*-nullcline, and a shifting of the fixed-point to the right (larger values of *V*). In other words, increasing values of *I_app_* make the *V*-nullcline “more cubic”. As for model 1, for low enough values of *I_app_* the fixed-point is a stable node (not shown) that transitions into a stable focus (Figs. 5-B1 and -B2). These small amplitude damped oscillations occur over a relatively small range of values of *V* by a mechanism similar to the one in model 1. However, in contrast to model 1, for model 2 the Hopf bifurcation is supercritical (in part due to the cubic-like shape of the *V*-nullcline). The small amplitude limit cycle generated as *I_app_* increases further (when the fixed-point loses stability) is stable (Fig. 5-B3) and its amplitude increases with increasing values of *I_app_* within some range (Fig. 5-B4). As mentioned above, in contrast to model 1, the onset of spikes is not described by the the two equations (for *V* and *r*) in model 2, and it requires a voltage threshold mechanism (Fig. 5-B4). The value of *V_th_* determines the maximal amplitude of the STOs.

In the following sections we investigate the consequences of the autonomous dynamics described above (Sections 3.1.1 and 3.1.2) for the subthreshold and suprathreshold responses of the *I_h_* + *I_Nap_* models to oscillatory input currents.

### 3.2 Models 1 and 2 exhibit different subthreshold gain dependencies with the input amplitude: supra- and sub-linear amplification of the voltage response

The subthreshold frequency preference properties of the *I_h_* + *I_Nap_* models 1 and 2 are captured by the impedance (Fig. 6-A1 and -B1), voltage envelope (Fig. 6-A2 and -B2) and phase (Fig. 6-A3 and -B3) profiles computed using eqs. (6). For the parameter values used, the two models exhibit both resonance and phasonance in the theta (4 – 12 Hz) frequency range. In [61, 70], we used a dynamic phase-plane analysis approach (e.g., Fig. 7) to explain the mechanisms of generation of resonance and phasonance for models with linear and parabolic *V*-nullclines.

**Figure 6:**
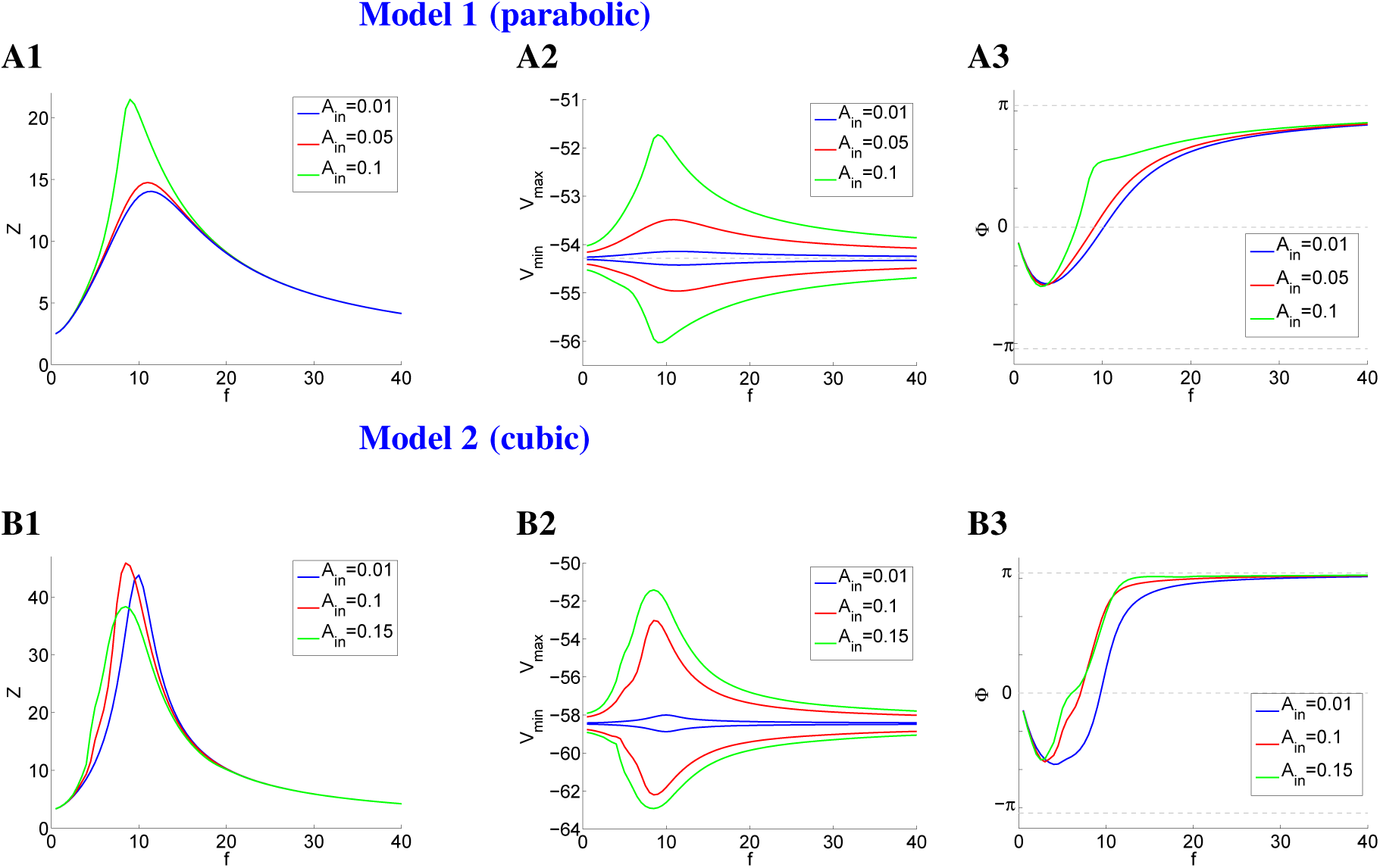
Impedance (left columns), voltage envelope (middle columns) and phase (right columns) profiles for the sinusoidally forced *I_h_*+*I_Nap_* models 1 (A) and 2 (B) for representative parameter values. **A.** *I_h_*+*I_Nap_* **model 1.** *We used the following parameter values: G_L_* = 0.5, *G_p_* = 0.5, *G_h_* = 1.5, *I_app_* = −2.5, *D* = 0. **B.** *I_h_*+*I_Nap_* **models 2.** *We used the following parameter values: G_L_* = 0.3, *G_p_* = 0.08, *G_h_* = 1.5, *I_app_* = 0.3, *D* = 0.

Briefly, the *V*-nullcline *N_V_* (*V*) (7) for the autonomous system (solid-red curves) moves cyclically following the sinusoidal input (5) in between the dashed-red curves obtained by substituting *I_app_* by *I_app_* ± *A_in_* in *N_V_* (*V*):

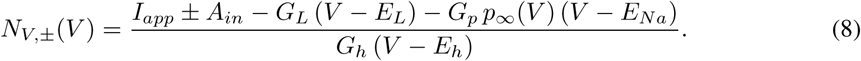

The moving *V*-nullcline

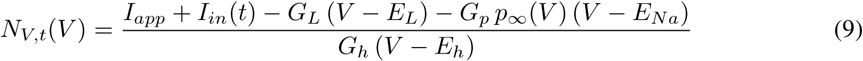
shifts down and rises on the input ascending and descending phases respectively, and reaches the minimum and maximum levels (dashed-red curves) at a quarter and three quarters of the input cycle respectively. We note that, technically, the *V*-nullcline does not move, but this motion is the visualization of a projection of the three-dimensional phase-space for *V, r* and *t* into the two-dimensional phase-plane for *V* and *r*.)

The response limit cycle (RLC) trajectories evolve in response to the motion of *N*_*V*,*t*_ with *f*-dependent speeds and directions, and therefore have *f*-dependent shapes. The values of *V_max_* and *V_min_* are given by the projections of the points with the maximum and minimum *V*-values on these limit cycle trajectories on the *V*-axis. In the limit of very low input frequency values (*f_in_* → 0) the RLC trajectory is slave to the motion of the *V*-nullcline, due to the slow motion of the latter. Therefore, the RLC motion occurs in the direction of the *r*-nullcline (e.g., *f_in_* = 0.5 in Figs. 7-A3 and -B3). In contrast, for very high input frequency values the RLC trajectory moves very slow as compared to the motion of the *N_V,t_*, and therefore the former has a very low amplitude and moves in a quasi horizontal direction (e.g., *f_in_* = 40 in Figs. 7-A3 and -B3). In the limit of *f_in_* → ∞, the RLC trajectory converges to the fixed-point for the autonomous system (not shown). For intermediate values of *f_in_* the RLC trajectory transitions in between these two extreme cases, by first rotating and becoming more elliptical, and then shrinking to a point as *A_in_* continues to increase. The resonant frequency correspond to the RLC trajectory that is able to reach the highest value *Z*(*f*) according to eq. (6). Under certain conditions (e.g., the number of input and output cycles coincide for each input frequency *f*) this corresponds to the optimal RLC trajectory that is able to reach the highest value of *V_max_* as compared to other input frequencies. From the point of view of the communication of the subthreshold frequency preference responses to the suprathreshold one, this is the measure that matters.

**Figure 7:**
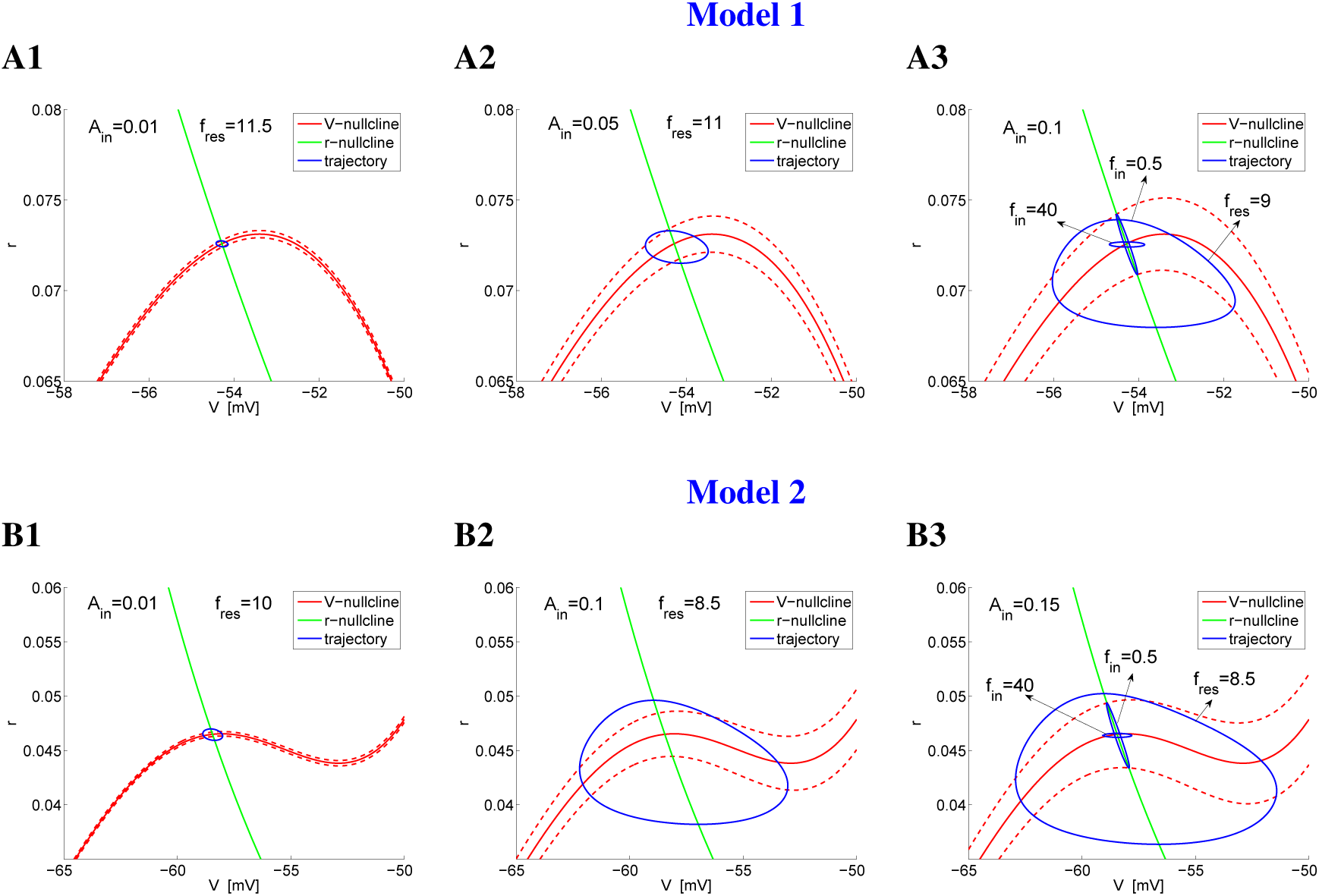
Phase-plane diagrams for the sinusoidally forced *I_h_*+*I_Nap_* models 1 (A) and 2 (B) for representative parameter values and the values of *A_in_* in Fig. 6. *The dashed-red curves are the V-nullclines displaced* ±*A_in_ units above and below the V-nullcline for the autonomous system (solid-red). They indicate the boundaries of the cyclic displacement of the V-nullcline as time progress due to the sinusoidal input. The V-nullcline reaches its lowest and highest levels (dashed-red curves) at a quarter and three quarters of each cycle respectively*. **A.** *I_h_*+*I_Nap_* **model 1.** *We used the following parameter values: G_L_* = 0.5, *G_p_* = 0.5, *G_h_* = 1.5, *I_app_* = −2.5, *D* = 0. **B.** *I_h_*+*I_Nap_* **models 2.** *We used the following parameter values: G_L_* = 0.3, *G_p_* = 0.08, *G_h_* = 1.5, *I_app_* = 0.3, *D* = 0.

As the input amplitude increases, the voltage response also increases (Fig. 7-A2 and -B2). For linear (or linearized) models this increase is proportional to the input amplitude *A_in_*, rendering the impedance independent of *A_in_*. For nonlinear models, the steady state responses to sinusoidal inputs violate at least one of the linearity principles: (i) coincidence of the output and input frequencies, (ii) proportionality between the output and the input, and (iii) symmetry of the output with respect to the equilibrium value around which the system is perturbed (resting potential). For the *I_h_* + *I_Nap_* models 1 and 2, (i) is satisfied, but not necessarily (ii) and (iii). In [61] we argued that the *I_h_* + *I_Nap_* model 1 exhibits a canard-related nonlinear amplification of the voltage response (Fig. 7-A1 and -A2) due to the ability of the resonant RLC trajectory to optimally follow the unstable branch of the parabolic-like *V*-nullcline for a significant amount of time (Fig. 6-A3). The combination of the parabolic-like shape of the *V*-nullcline and the time scale separation between the participating variables allows the resonant RLC trajectory to reach higher values of *V* as compared to the resonant RLC trajectory for the corresponding linearized system (not shown). (The latter would have a more rounded shape as in Fig. 6-A2). We refer to this type of nonlinear amplification of the voltage response as supra-linear.

In contrast to the *I_h_* + *I_Nap_* model 1, the *I_h_* + *I_Nap_* model 2 exhibits a sub-linear amplification of the voltage response (Fig. 6-B1 and -B2), which implies a negative gain. Specifically, for low enough values of *A_in_* (e.g., *A_in_* = 0.1 in Fig. 6-B) the amplification is supra linear, but it becomes sub-linear as *A_in_* increases further (e.g., *A_in_* = 0.15 in Fig. 6-B). The impedance profiles for *A_in_* = 0.15 is below the ones for *A_in_* = 0.1 and *A_in_* = 0.01. We use the latter case as representing the linear behavior due to the low input amplitude and the symmetric properties of the envelope amplitude diagrams (Fig. 7-A1 and A2).

The initial supra-linear amplification occurs when the RLC trajectories, in particular the resonant ones, move around the parabolic-like part of the cubic-like *V*-nullcline (Fig. 7-B2), by a mechanisms similar to the one described above. However, this mechanism is disrupted as *A_in_* increases further and the RLC trajectories enter the voltage regime where the cubic-like *V*-nullcline “bends up”, thus changing the effect the vector field has on the shape of the RLC trajectories.

In contrast to subthreshold resonance, subthreshold phasonance does not show a significant qualitative difference in the monotonic properties between the models 1 and 2 (Fig. 7-A3 and -B3). This is consistent with previous findings showing that resonance and phasonance are related, but different phenomena [59, 61].

These results persist for a larger range of values of *τ_r_* (not shown). The differences between the two models are more pronounced for larger than for smaller values of *τ_r_*. For model 2 and smaller values of *τ_r_* (e.g., *τ_r_* = 40) the impedance profile still decreases as *A_in_* increases, but it may require larger values of *A_in_* for the impedance profile to decrease below the linear impedance.

Since generating spiking responses involves increasing the values of *A_in_* above these used to obtain subthreshold resonance, the results above suggest that the communication of the subthreshold frequency preferences to oscillatory inputs to the suprathreshold regime, when it happens, will have different properties between the *I_h_* + *I_Nap_* models 1 and 2. We address this in the following sections.

### 3.3 Suprathreshold spiking resonance: evoked, output and phase responses

Below we examine the suprathreshold frequency preference response of the *I_h_* + *I_Nap_* models 1 and 2 to sinusoidal inputs of the form (5). We primarily address two issues: (i) whether and how the subthreshold resonant properties are communicated to the suprathreshold regime, and (ii) what are the differences and similarities between the suprathreshold responses to oscillatory inputs between the two *I_h_* + *I_Nap_* models.

We will focus on three different types of spiking resonances that occur depending on whether one focuses on (i) the input frequencies (what input frequencies are able to generate spikes), (ii) the output frequencies (regardless of the input frequencies that generate the response), and (iii) the phase-shift of the spiking output with respect to the input peak (spiking phase). While for subthreshold resonance, one can identify a single number (the resonant frequency *f_res_*) as defining the preferred frequency response, for the spiking resonances it is often more convenient to identify resonant frequency bands Δ*f_res_* for which the models exhibit preferred spiking responses. Here we focus on Δ*f_res_* in the theta frequency regime, which is the frequency band at which the models we use exhibit subthreshold resonance.

*Evoked spiking resonance* occurs when spiking is generated only for input frequencies in an intermediate (resonant) frequency band Δ*f*_*res*,*ev*_. *Output spiking resonance* occurs when the output frequency primarily belongs to an intermediate (resonant) frequency band Δ*f*_*res*,*out*_, regardless of the input frequency that generated the response. *Spiking phasonance* occurs when the phase-shift (phase) Φ*_spk_* between the output spike and the input peak vanishes at a non-zero input frequency. We provide more details below in our discussion of the specific cases.

The definition of spiking phasonance is the natural extension of subthreshold phasonance, using (6), to the suprathreshold regime. It is rather restrictive for two reasons. First, it assumes that a single spike is produced for each input cycle. However, Φ*_spk_* can be computed when more than one spike are produced by a single input cycle. Second, it does not take into account the cases where Φ*_spk_* does not vanish for any value of the input frequency, but it is minimized for a given input frequency. We will not consider this situation in this paper.

The mechanisms that govern the transition from the subthreshold to the spiking resonances depend on the interplay of the sinusoidal input and the autonomous spiking dynamics described in Section 3.1.2. The latter, in turn, depends on the subthreshold dynamics and the spiking mechanism for the two models.

We leave out of this study the analysis of other types of preferred frequency responses to oscillatory inputs such as the firing rate resonance [2] where the neuron’s firing rate (e.g., see [74]) response *R_out_* (*f*) measured in terms of the signal gain *A*(*f*) = *R_out_*(*f*)/*A_in_* peaks at a preferred input frequency.

### 3.4 Suprathreshold spiking resonance: similarities and differences between the *I_h_* + *I_Nap_* models 1 and 2

#### 3.4.1 Evoked and output spiking resonance are inherited from the neuronal subthreshold resonant properties for small values of *A_in_* in models 1 and 2

For low enough values of *A_in_* subthreshold resonance is communicated to the suprathreshold regime to produce both evoked and output spiking resonance (Figs. 8-A1 and 10-A1) in input frequency bands Δ*f*_*res*,*ev*_ and Δ*f*_*res*,*out*_) around 10 Hz in both models. The existence of these resonances is a direct consequence of the fact that a small enough increase in *A_in_* above the subthreshold values causes spiking both within a limited range of output frequencies and for only a small range of input frequencies around the subthreshold frequency band. The output/input frequency patterns are different between the two models: 2:1 for the model 1 (Figs. 8-A1) and (mostly) 1:1 for the model 2 (Fig. 10-A1). The differences in the output/input patterns reflect the geometric differences between the two *I_h_* + *I_Nap_* models.

**Figure 8:**
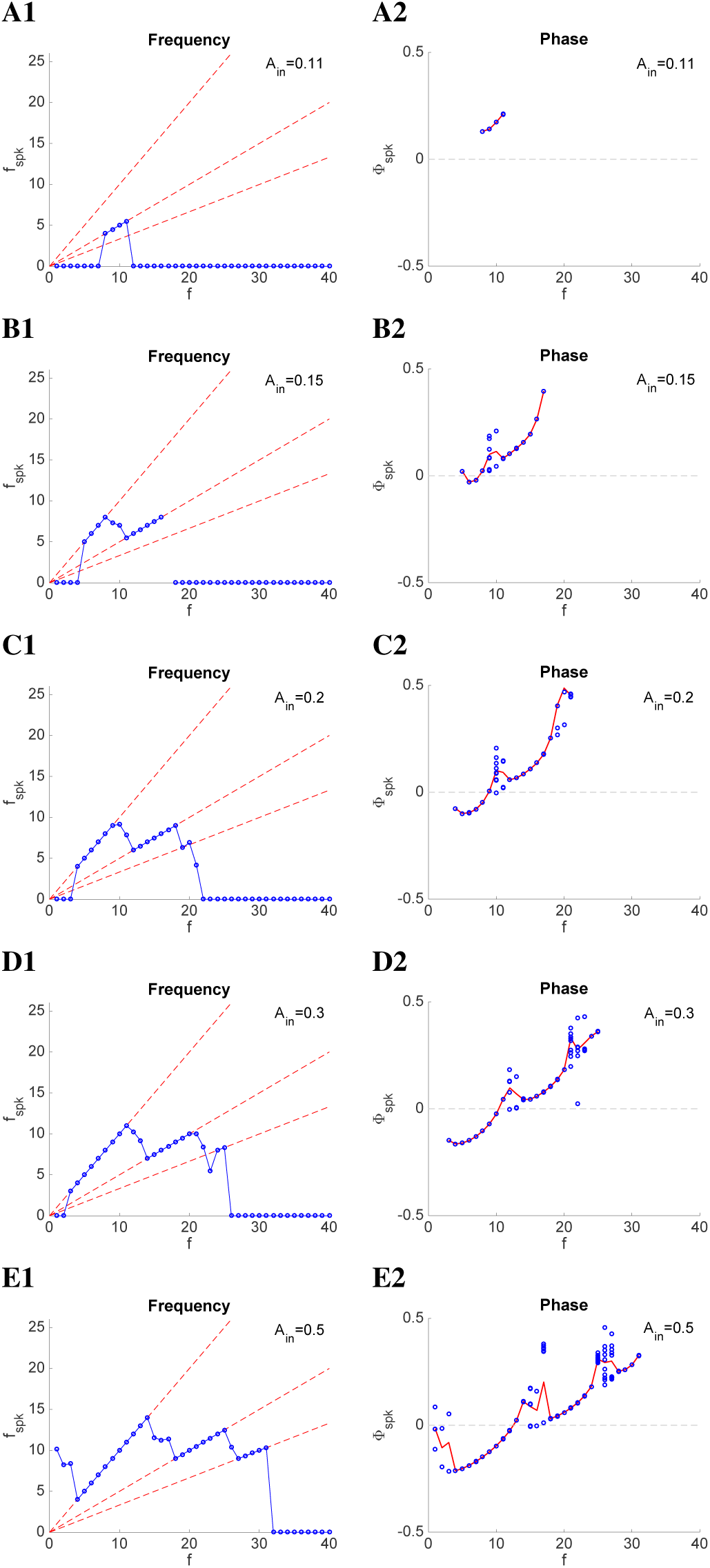
Suprathreshold response of the *I_h_*+*I_Nap_* model 1 to sinusoidal inputs for representative parameter values (*as in Figs. 7-A*). **Left panels**: *Spike-frequency diagrams. The output spike frequency f_spk_ is the normalized inverse of the average length of the interspike intervals (Hz). The dashed-red lines (from top to bottom) indicate the 1:1, 2:1, and 3:1 output spikes versus input cycle patterns, respectively*. **Right panels:** *Spike-phase diagrams. The output spike phase* Φ*_spk_ (blue dots) was computed as the difference between the output spike-time and the closest input peaktime normalized by the cycle length*. Φ*_spk_* = 0 *for spikes at the input peak and* Φ*_spk_* = ±0.5 *for spikes at the immediate prior and posterior input troughs. The red line indicates the average* Φ*_spk_ for each input frequency. We used the following parameter values: G_L_* = 0.5, *G_p_* = 0.5, *G_h_* = 1.5, *I_app_* = −2.5, *D* = 0, *V_th_* = −45, *V_rst_* = −75, *r_rst_* = 0.

In model 1, spiking resonance is created by an extension of the canard-like mechanism described above (Fig. 7-A) [61]. Figure 9-A illustrates this for *A_in_* = 0.11 (the value used in Fig. 8-A, just above the value used in Fig. 7-A: *A_in_* = 0.1). For input frequencies below and above Δ*f_res_* (Figs. 9-A1 and -A4 resp.), the limit cycle trajectory moves around the *N*_*V*,*t*_ knee and crosses it when *N*_*V*,*t*_ is raising (descending phase), thus creating STOs. The differences in the shapes of these small amplitude limit cycles reflect the dependence of the RLC trajectory on the interaction between the input frequency time scale and intrinsic dynamics structure (nonlinearities and time scale separation) described in Section ??.

**Figure 9:**
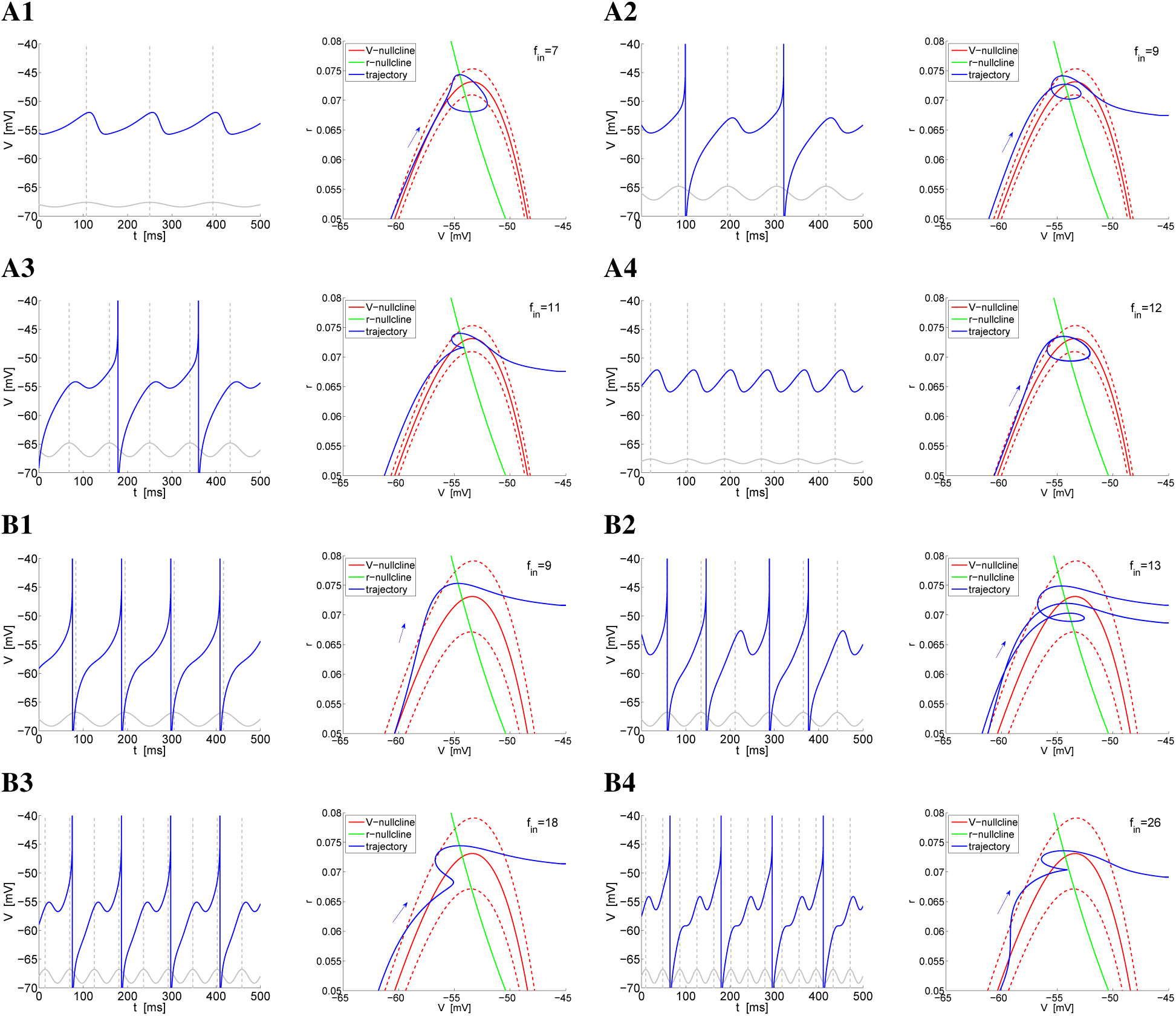
Suprathreshold response of the *I_h_*+*I_Nap_* model 1 to sinusoidal inputs for representative parameter values and *A_in_* = 0.3. *Voltage traces (left panels) and phase-plane diagrams (right panels) for representative values of A_in_ and f_in_*. **A.** *A_in_*, = 0.11. **B.** *A_in_*, = 0.3. *Parameter values are as in Figs. 7-A and Fig. 8-D*. **Left panels:** *The solid-gray curves are caricatures of the sinusoidal inputs. The dashed-gray vertical lines at the peaks of the sinusoidal inputs indicate* Φ*_spk_* = 0 *(zero phase-shift)*. **Right panels:** *The dashed-red curves are the V-nullclines displace* ±*A_in_ units above and below the V-nullcline for the autonomous system (solid-red). They indicate the boundaries of the cyclic displacement of the V-nullcline as time progress due to the sinusoidal input. The V-nullcline reaches its lowest and highest levels (dashed-red curves) at a quarter and three quarters of each cycle respectively.The arrow indicates the direction of motion of the trajectory from its reset point to the spiking regime. We used the following parameter values: G_L_* = 0.5, *G_p_* = 0.5, *G_h_* = 1.5, *I_app_* = −2.5, *D* = 0, *V_th_* = −45, *V_rst_* = −75, *r_rst_* = 0.

As *f_in_* increases above the values in Figs. 9-A1 *N*_*V*,*t*_ moves faster and opens a window of opportunity for the RLC trajectory to escape the subthreshold regime and produce a spike. Figs. 9-A2 and -A3 (left) shows the 2:1 MMO patterns for *f_in_* = 9 and *f_in_* = 11 respectively. The phase-plane diagrams (right) show the corresponding trajectories initially at the reset values (*V_rst_* and *r_rst_*) during the descending phase of the input. The loops in the phase-plane diagrams reflect the STOs in the left panels.

The trajectory first moves along the left branch of *N*_*V*,*t*_ as it raises from its minimum level (input peak) towards its maximum level (input trough), and then shifts down again. The trajectory reaches the knee when the trajectory is near its baseline level and evolves around the knee as it continues to shift down, to reach its minimum level. The STO is created because the trajectory is moving around the right branch of the *N*_*V*,*t*_ while the latter is beginning to raise, and therefore they intersect.This intersection occurs at a higher level (“earlier”) as compared to *f_in_* = 7, therefore the STO in Fig. 9-A2 has a smaller amplitude than in Fig. 9-A1. The STO trajectory crosses the left branch of *N*_*V*,*t*_ and reaches the left branch when the trajectory is near its maximum level, then it moves around the *N*_*V*,*t*_ knee as it shifts down. In contrast to the previous input cycle, when the trajectory reaches the right branch, the *N*_*V*,*t*_ is shifting down, therefore the distance between the two is large enough to allow the trajectory to move away from the vicinity of *N*_*V*,*t*_ towards the spiking regime before *N*_*V*,*t*_ raises back. As *f_in_* increases further (*f_in_* = 11; Fig. 9-A3) the speed of *N*_*V*,*t*_ increases, causing the trajectory to cross the *N*_*V*,*t*_ at lower level (as compared to *f_in_* = 9) during the first input cycle, therefore creating a STO with a smaller amplitude.

When *f_in_* increases to *f_in_* = 12 (Fig. 9-A4) *N*_*V*,*t*_ moves faster than for *f_in_* = 11 and the limit cycle trajectory looses the ability to generate a STO on the left branch of *N*_*V*,*t*_. Instead, it continues to move and crosses the *N*_*V*,*t*_ on its right branch, when it is shifting down, thus creating a STO with a larger amplitude than for *f_in_* = 11, but there is no spiking.

In model 2, spiking resonance is created by a different mechanism that for model 1 (Fig. 11-A), which involves the interplay of the cubic-like subthreshold dynamics and a voltage threshold for spike generation.

**Figure 10:**
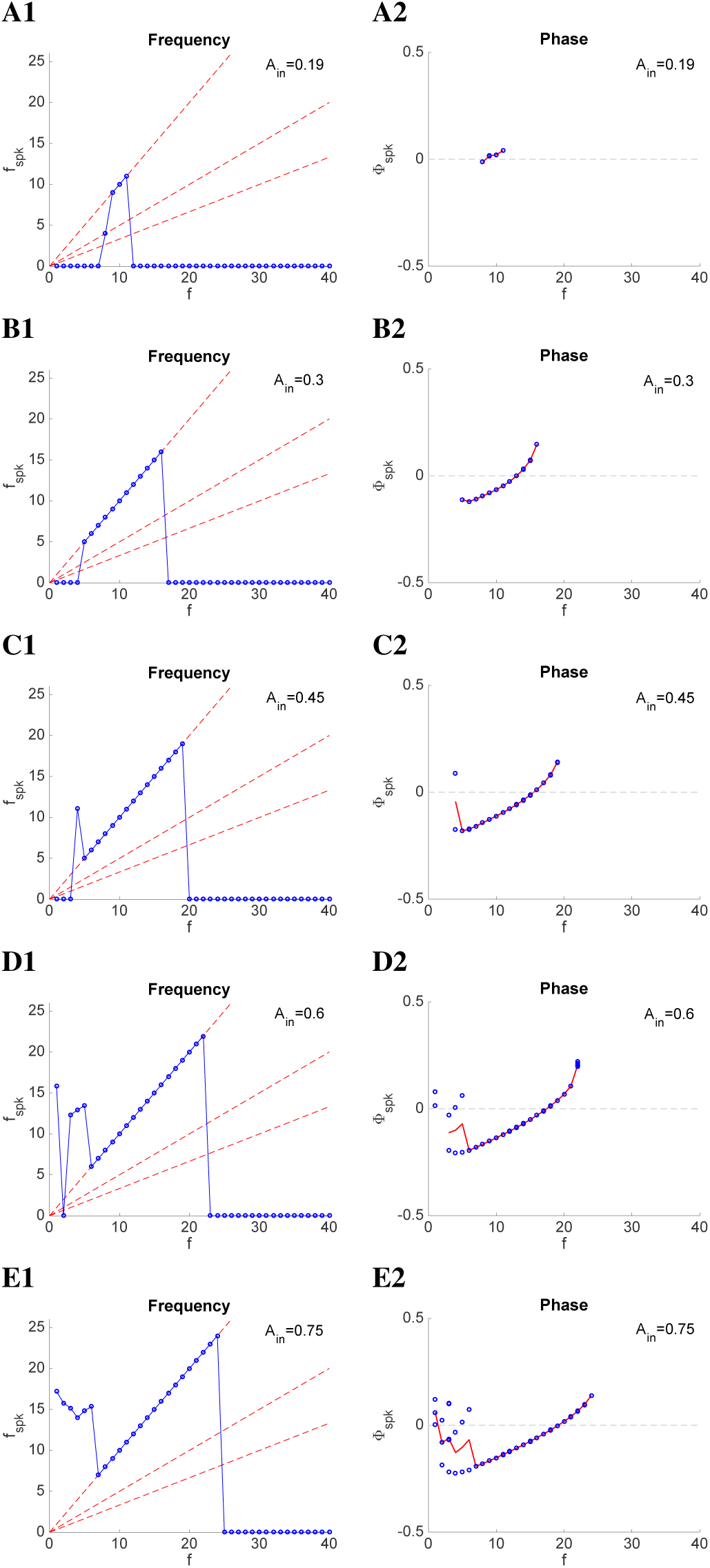
Suprathreshold response of the *I_h_*+*I_Nap_* model 2 to sinusoidal inputs for representative parameter values (*as in Figs. 7-B*). **Left panels: Spike-frequency diagrams. The output spike frequency** *f_spk_* **is the normalized inverse of the average length of the interspike intervals (Hz). The dashed-red lines (from top to bottom) indicate the 1:1, 2:1, and 3:1 output spikes versus input cycle patterns, respectively. Right panels: Spike-phase diagrams. The output spike phase** Φ,*_spk_* **(blue dots) was computed as the difference between the output spike-time and the closest input peak-time normalized by the cycle length.** Φ*_spk_* = 0 **for spikes at the input peak and** Φ*_spk_* = ±0.5. **for spikes at the immediate prior and posterior input troughs. The red line indicates the average** Φ*_spk_* **for each input frequency. We used the following parameter values:** *G_L_* = 0.3, *G_p_* = 0.08, *G_h_* = 1.5, *I_app_* = 0.3, *D* = 0, *V_th_* = −51, *V_rst_* = −75, *r_rst_* = 0.

**Figure 11:**
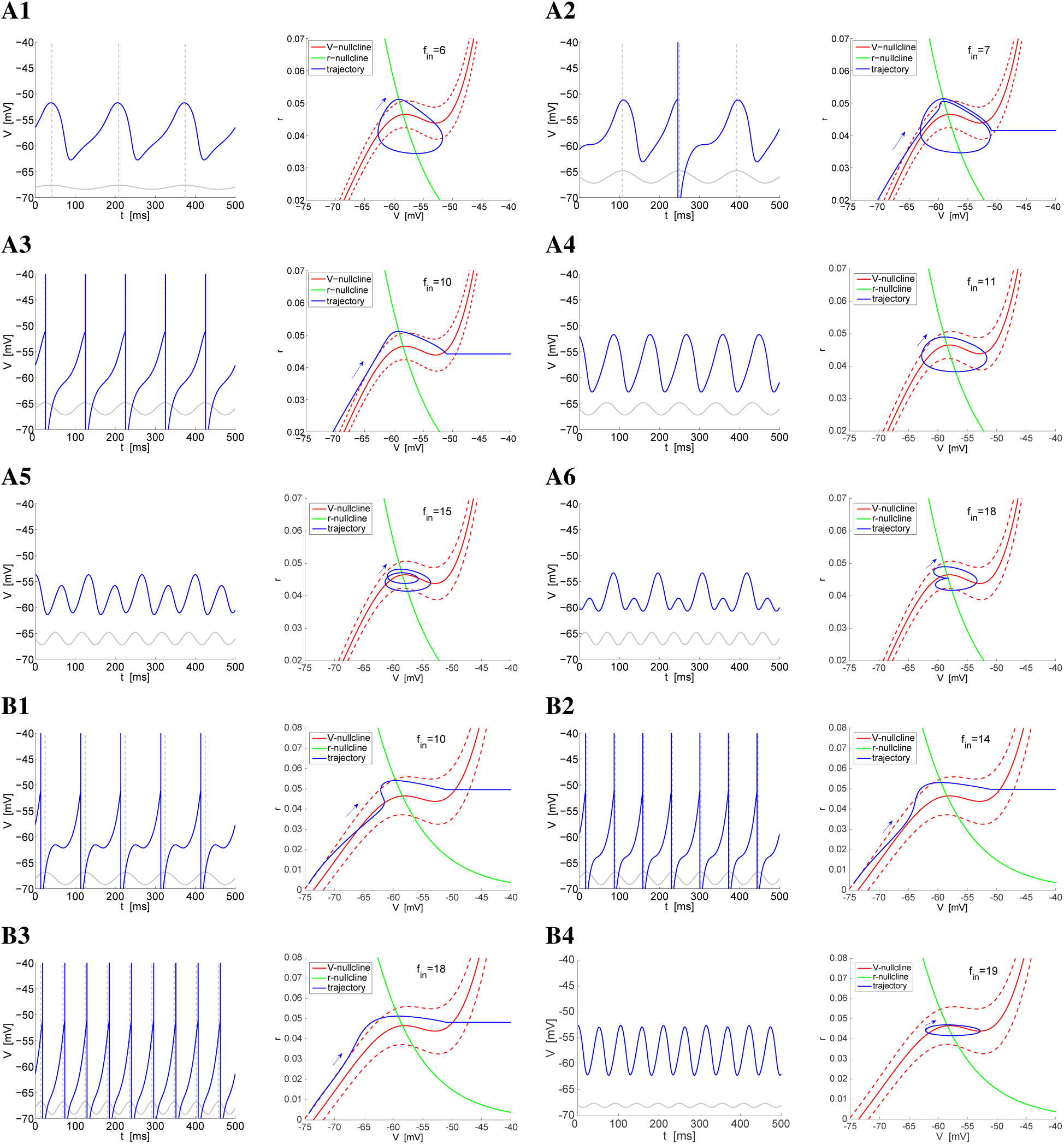
Suprathreshold response of the *I_h_*+*I_Nap_* model 2 to sinusoidal inputs for representative parameter values. *Voltage traces (left panels) and phase-plane diagrams (right panels) for representative values of A_in_ and f_in_. Parameter values are as in Figs. 7-B and Fig. 8-C*. **A.** *A_in_* = 0.19. **B.** *A_in_* = 0.45. **Left panels:** *The solid-gray curves are caricatures of the sinusoidal inputs. The dashed-gray vertical lines at the peaks of the sinusoidal inputs indicate* Φ*_spk_* = 0 *(zero phase-shift)*. **Right panels:** *The dashed-red curves are the V-nullclines displace* ±*A_in_ units above and below the V-nullcline for the autonomous system (solid-red). They indicate the boundaries of the cyclic displacement of the V-nullcline as time progress due to the sinusoidal input. The V-nullcline reaches its lowest and highest levels (dashed-red curves) at a quarter and three quarters of each cycle respectively.The arrow indicates the direction of motion of the trajectory from its reset point to the spiking regime. We used the following parameter values: G_L_* = 0.3, *G_p_* = 0.08, *G_h_* = 1.5, *I_app_* = 0.3, *D* = 0, *V_th_* = −51, *V_rst_* = −75, *r_rst_* = 0.

For input frequencies below and above Δ*f_res_* (Figs. 11-A1 and -A4 to -A6 resp.), the RLC trajectories never cross the *V_th_* line. As for the model 1, the differences in the shapes of these RLCs reflect the dependence of the model response on the interaction between the input frequency time scale that causes *N*_*V*,*t*_ to raise and shift down, and the intrinsic cubic-like dynamic structure (see Section ??).

Although there is a time scale separation between the participating variables (the value of *τ_r_* for the models 1 and 2 are the same), the STOs are not of relaxation type as it would occur for the “classical” cubic-like models such as the FitzHugh-Nagumo model [75, 76] (see also [77]), but more rounded due to the small difference between the maximum and the minimum of the *V*-nullcline as explained earlier.

For low values of *f_in_*, just outside Δ*f_res_* (e.g., *f_in_* = 6; Fig. 11-A1) the trajectory moves up along the *V*-nullcline as it raises towards its maximum level. The response trajectory reaches the upper knee roughly at the same time as *N*_*V*,*t*_ reaches its maximum level and moves down in a vicinity of the right branch accompanying *N*_*V*,*t*_ as it shifts down, then crossing roughly when *N*_*V*,*t*_ reaches its minimum level at a value of *V* < *V_th_*.

For high enough values *f_in_* just outside Δ*f_res_* (e.g., *f_in_* = 11; Fig. 11-A4) *N*_*V*,*t*_ moves faster than for *f_in_* = 6 and reaches its maximum level while the response trajectory is moving along the left branch, but relatively far away from the upper knee. Because of the larger distance between the trajectory and the fixed-point, the response trajectory moves in a more horizontal direction in the vicinity of the middle branch, but reaches values of *V* < *V_th_*.

For values of *f_in_* within Δ*f_res_* the response trajectory reaches large enough values of *V* large enough to reach (and cross) the *V_th_* line to produce spikes. The transition from STO to spiking responses includes a very small range of input frequencies for which the system exhibits a MMO response. This results from a combination of the dynamics of the system for smaller (e.g., *f_in_* = 6) and larger (e.g., *f_in_* = 8) input frequencies. Specifically, during one input cycle, after a spike has occurred, the response trajectory follows *N*_*V*,*t*_ as it raises from its minimum level and reaches the upper knee roughly at the same time as *N*_*V*,*t*_ reaches its maximum level. The response trajectory slows down as a consequence of its proximity to the fixed-point while *N*_*V*,*t*_ shifts down. The increasing distance between the two causes the response trajectory to intersect the right side of the lower knee at a value of *V* < *V_th_*. During the second cycle, the response trajectory crosses the left branch when *N*_*V*,*t*_ is already raising above its baseline, thus allowing the response trajectory to reach a higher value of *V* and cross the *V_th_* line.

#### 3.4.2 Evoked theta spiking resonance vanishes for larger values of *A*in**: evoked broadband and low-pass filters

Both models 1 and 2 exhibit evoked spiking resonance for low enough values of *A_in_* above the subthreshold level because only a small portion of the upper envelope of the voltage response, around the subthreshold resonance peak (Figs. 6-A2 and -B2), raises above threshold. As *A_in_* increases further, the range of input frequencies that is able to evoke a spiking response expands (Figs. 8-B1 to -E1 and 10-B1 to -E1), eventually creating an evoked spiking low-pass filter (Figs. 8-E1 and 10-E1), where all input frequencies below some value produce a spiking response. While for intermediate values of *A_in_* (e.g., Figs. 8-B1 to -D1 and 10-B1 and -C1) the input frequency band that is able to evoke a spiking response is bounded from both below and above, it is too broad and exceeds the theta frequency band.

The mechanisms that prevented spiking for input frequencies outside Δ*f_res_* in Figs. 8-A1 and -A4 (model 1) and 10-A1 and -A4 (model 2) are dependent on balances between the speed of motion of *N*_*V*,*t*_ (a input frequency time scale) and the input amplitude *A_in_* that causes the trajectory to cross the right branch before it is able to escape the subthreshold regime (model 1) or cross the *V_th_* line (model 2). Increasing values of *A_in_* disrupt this balance, thus generating spiking for a broader range of input frequencies.

#### 3.4.3 Spiking phasonance is created as *A_in_* increases and persists for larger values of *A_in_*

Spiking phasonance (Figs. 8-B2 to -E2 an 10-B2 to -E2) occurs when the neuron spikes at the peak of the input cycle. This phenomenon also requires a balance between the speed of motion of *N*_*V*,*t*_ and the input amplitude *A_in_* so that the trajectory is neither too fast nor too slow as compared to the dynamics of *N*_*V*,*t*_ and is able to reache the spiking regime exactly at the same time as *N*_*V*,*t*_ reaches its minimum level (input peak). If *A_in_* is too small, then the subthreshold response may be synchronized in phase with the input oscillations, but no spikes are produced. Clearly, the mechanism of generation of spiking phasonance depends on the mechanisms of spike generation and trajectory reset after a spike has occurred. We consider other scenarios later in the paper.

#### 3.4.4 Model 1 exhibits theta output resonance for intermediate values of *A_in_*

For small enough suprathreshold values of *A_in_* the output spiking frequency increases with increasing values of the input frequency *f* (Fig. 8-A1) and both the evoked and output frequency bands are within the theta range. As *A_in_* increases within some range (Figs. 8-B1, -C1 and -D1), the output frequency band remains within the theta range, while the evoked frequency band increases beyond theta frequencies. The spiking frequency patterns (left panels) are non-monotonic functions of *f* showing the existence complex patterns including the 2:1 and 3:1 ones in addition to the 1:1 for lower values of *f*. These patterns are generated by cycle skipping mechanisms (Figs. 9-B, left panels) as *f* increases beyond the theta range.

Because of the higher value of *A_in_* the *V*-nullclines in Fig. 9-B reach higher and lower levels as they raise and shift down, respectively, following the dynamics of the sinusoidal input as compared to the *V*-nullclines in Fig. 9-A. This allows the response trajectory for the 1:1 pattern in Fig. 9-B1 to reach the spiking region of the phase-plane while the *V*-nullcline is near its minimum level, and therefore it produces a spike without “interferences”. More specifically, the response trajectory moves along *N*_*V*,*t*_ as it raises during the descending phase of the input. During the ascending phase, while *N*_*V*,*t*_ shifts down, the response trajectory moves around the knee of the *N*_*V*,*t*_ and produces a spike when the *V*-nullcline is close to its minimum level.

As *f* increases the response trajectory evolves slower. For *f* above the theta range (Figs. 9-B3) *N*_*V*,*t*_ completes a full cycle (returns to the solid-red baseline) when the response trajectory is still moving along it, but still didn’t reach the knee. The response trajectory reaches the knee when *N*_*V*,*t*_ is raising from its minimum level on the subsequent cycle. As this happens, the response trajectory is “caught inside” the *V*-nullcline and therefore is forced to reverse direction and move along the *N*_*V*,*t*_ as it continues to raise, giving rise to the bump STO. Spiking occurs within this second cycle when after the *V*-nullcline shifts down.

A similar mechanism is responsible for the generation of theta output patterns in Fig. 9-B4. However, in these 3:1 patterns spiking occurs close to the trough of the input signal instead of its peak as for the 1:1 and 2:1 patterns in Figs. 9-B1 to -B3. For the 3:1 patterns the onset of spikes occurs as the response trajectory is able to move past the knee of the *V*-nullcline when the latter is raising, without being “caught inside”. Spiking occurs later in the cycle when the *V*-nullcline is at a higher level than the response trajectory. For this to happen it is crucial that the speed of the response trajectory is high enough to overcome the motion of the *V*-nullcline. As *f* increases further, the speed of the response trajectory is lower, and spiking is no longer produced. Instead, the response trajectory moves around the knee of the *V*-nullcline (not shown) as in Fig. 9-A4.

#### 3.4.5 Model 2 exhibits a broad-band output response for intermediate values of *A_in_*

Similar to model 1, for small enough suprathreshold values of *A_in_* the output spiking frequency increases with increasing values of the input frequency *f* (Fig. 8-A1) and both the evoked and output frequency bands are within the theta range. In contrast to model 1, the response patterns of model 2 are 1:1. As *A_in_* increases (Figs. 10-B1 to -D1) the output spiking patterns continue to be 1:1 and the output spiking frequency increases linearly with the input frequency. Therefore, when the evoked frequency band is outside the theta range so does the output frequency band.

The qualitative differences between the output patterns in the two models are due to the differences in their dynamic structures, particularly both the shapes of the *V*-nullclines and the underlying vector fields, and the spiking mechanisms. In model 2 (Fig. 11-B1 to -B3), the response trajectory moves in a vicinity of the *V*-nullcline, first as the *V*-nullcline raises (descending input phase) and then as it shifts back down (ascending input phase). Spiking is produced as long as the trajectory is fast enough to reach threshold without intersecting the *V*-nullcline. Otherwise, response STOs are produced (Fig. 11-B4).

Model 2 does not exhibit theta output spiking resonance because the cubic-like dynamic structure does not admit the type of 2:1 and 3:1 MMO response patterns displayed by model 1. In contrast to model 1, the response trajectory for model 2 moves along the upper dotted *V*-nullcline during the initial portion of the cycle (compare Figs. 11-B3 and 9-B3) as the *V*-nullcline raises. When the *V*-nullcline shifts down, the response trajectory is away from the region of slow motion and it moves faster towards the spiking regime than the response trajectory in model 1 (Fig. 9-B3) when it arrives at the vicinity of the parabolic-like nullcline during the STO response cycle.

One could in principle think that the fact that the baseline *V*-nullcline in model 1 is higher than the baseline *V*-nullcline in model 2 plays a role in determining the response patterns in the two models. Specifically, it would be plausible to think that the response trajectory for model 2 is able to reach threshold without displaying MMO response patterns as does model 1 simply because it has to move along a shorter distance than the response trajectory for model 1. In order to rule out this possibility we modified models 1 and 2 in such a way that the baseline *V*-nullcline for model 1 is lower (Fig. 12) and th baseline *V*-nullcline for model 2 is higher (13). Fig. 12 shows that model 1 displays the same type of MMO response patterns as in Fig. 9-B and Figs. 13-A1 to -A3 shows that MMO response patterns are absent for model 2 for comparable input frequencies as in Fig. 11-B. Figs. 13-A4 and -A5 show MMO patterns for higher frequencies reflecting the fact that the transition from response spiking to absence of spikes is not abrupt as in Fig. 10. These MMO response patterns are generated by a different mechanism than these more model 1 (e.g., Fig. 12) where the STO portions of the response trajectory move around the cubic-like *V*-nullcline.

**Figure 12:**
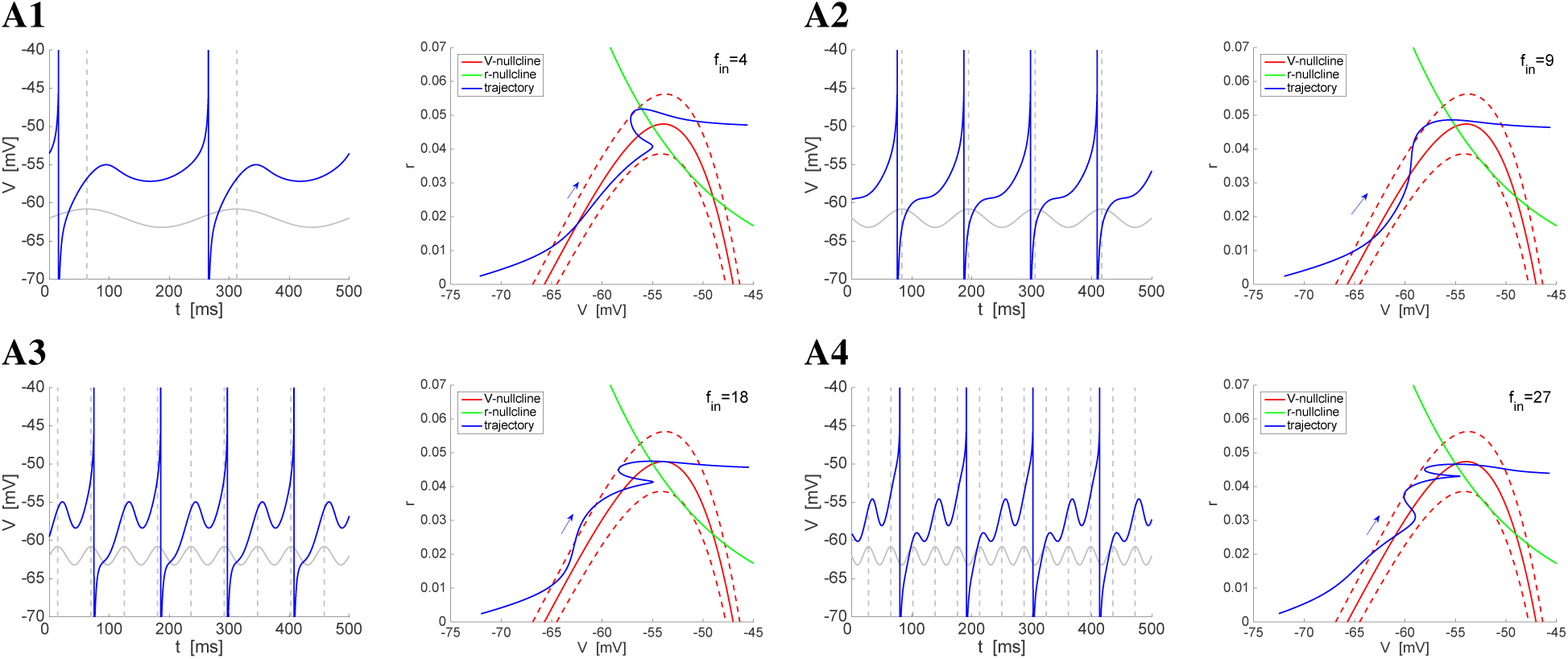
Suprathreshold response of the *I_h_*+*I_Nap_* model 1 to sinusoidal inputs for representative parameter values and *A_in_* = 0.45. *Voltage traces (left panels) and phase-plane diagrams (right panels) for representative values of f_in_. Parameter values are as in Figs. 8 and 9 (and Figs. 7-A and Fig. 8-D) with two exceptions: The r-nullcline was shifted to the left to accommodate the change in I_app_ that shifted the V-nullcline down*. **Left panels:** *The solid-gray curves are caricatures of the sinusoidal inputs. The dashed-gray vertical lines at the peaks of the sinusoidal inputs indicate* Φ*_spk_* = 0 *(zero phase-shift)*. **Right panels:** *The dashed-red curves are the V-nullclines displace* ±*A_in_ units above and below the V-nullcline for the autonomous system (solid-red). They indicate the boundaries of the cyclic displacement of the V-nullcline as time progress due to the sinusoidal input. The V-nullcline reaches its lowest and highest levels (dashed-red curves) at a quarter and three quarters of each cycle respectively.The arrow indicates the direction of motion of the trajectory from its reset point to the spiking regime. We used the following parameter values: G_L_* = 0.5, *G_p_* = 0.5, *G_h_* = 1.5, *I_app_* = −1.2, *D* = 0, *V_th_* = −45, *V_rst_* = −75, *r_rst_* = 0.

#### 3.4.6 Theta output resonance vanishes for higher values of *A_in_* for model 1

As *A_in_* increases further the output frequency band for model 1 increases beyond the theta regime (Figs. 8-E) as the result of the increase in the length of the 1:1 response branch. The increase in *A_in_* causes the *V*-nullcline to move further away from the baseline *V*-nullcline, thus allowing the generation of spikes for a larger range of input frequencies without the response trajectories being “caught” by the *V*-nullcline on its way back up (descending phase) and being forced to produce STOs. In spite of the fact that theta output resonance is lost, the evoked spiking response remains in a relatively bounded output frequency band.

#### 3.4.7 Spiking phasonance is not necessarily inherited from subthreshold phasonance for small values of *A_in_*

Fig. 8-A2 shows that the existence of subthreshold phasonance does not necessarily imply the generation of spiking phasonance even for small values of *A_in_*. For this to occur at least two conditions need to be satisfied: (i) *f_phas_* has to be close enough to *f_res_*, and (ii) the onset of spikes in response to the sinusoidal inputs has to be fast enough. In the limit, both *f_res_* = *f_phas_* and the onset of spikes has to be instantaneous. From our previous discussion, the onset of spikes is faster for model 2 than for model 1 and therefore it is not surprising that the latter exhibits spiking phasonance (Fig. 10-A2), while the former does not.

#### 3.4.8 Model 1 exhibits theta spiking phasonance for higher values of *A_in_*

As *A_in_* increases spiking occurs at earlier phases for the 1:1 patterns in model 1, and therefore it exhibits spiking phasonance at theta frequencies (Fig. 8-B2 to -E2, right). The irregular patterns (in between pure 1:1 and 2:1) show a bimodal Φ*_spk_* distribution with the lower values of Φ*_spk_* close to phasonance. This persists for higher values of *A_in_* within some range (Fig. 8-B2 to -D2, right). Above this range (Fig. 8-E2, right) model 1 exhibits phasonance at theta frequencies for the regular patterns and above the theta frequency range for the irregular patterns.

#### 3.4.9 The phasonant frequency in model 2 increases with increasing values of *A_in_* and exceeds the theta frequency range for high enough values of *A_in_*

This is shown in Figs. 11 (right). The patterns are more regular than for model 1, except for the combination of lower frequencies and higher values of *A_in_* that produce burst-like patterns (Figs. 11-D2 and -E2, right). The monotonic dependence of *f_phas_* with *A_in_* is due to the fact that as *A_in_* increases, spiking occurs earlier in the cycle for for a given input frequency because for these frequencies *V* crosses threshold at lower values. For spiking to occur at the input peak time, the input frequency has to be higher.

#### 3.4.10 The response patterns for both models 1 and 2 do not qualitatively change for higher values of *V_rst_* and *r_rst_*

Here we investigate whether the spiking patterns obtained in the previous sections for models 1 and 2 and the qualitative differences between the patterns for the two models depend on the specific reset values *V_rst_* and *r_rst_*. In Figs. 8 to 13 (*V_rst_, r_rst_*) is to the left of the stable fixed-point in the respective phase-plane diagrams and *r_rst_* = 0. The response patterns involve the evolution of the response trajectories along the corresponding slow manifolds until they reach the region of parabolic- or cubic-like nonlinearities according to the model type. Here we consider models 1 and 2 with the same parameter values as before (Figs. 8, 10 and 12 for model 1 and Figs. 9, 11 and 13 for model 2), except for (*V_rst_, r_rst_*), which is to the right of the fixed-point in the phase plane and within the region of the corresponding nonlinearities as in Fig. 3.

**Figure 13:**
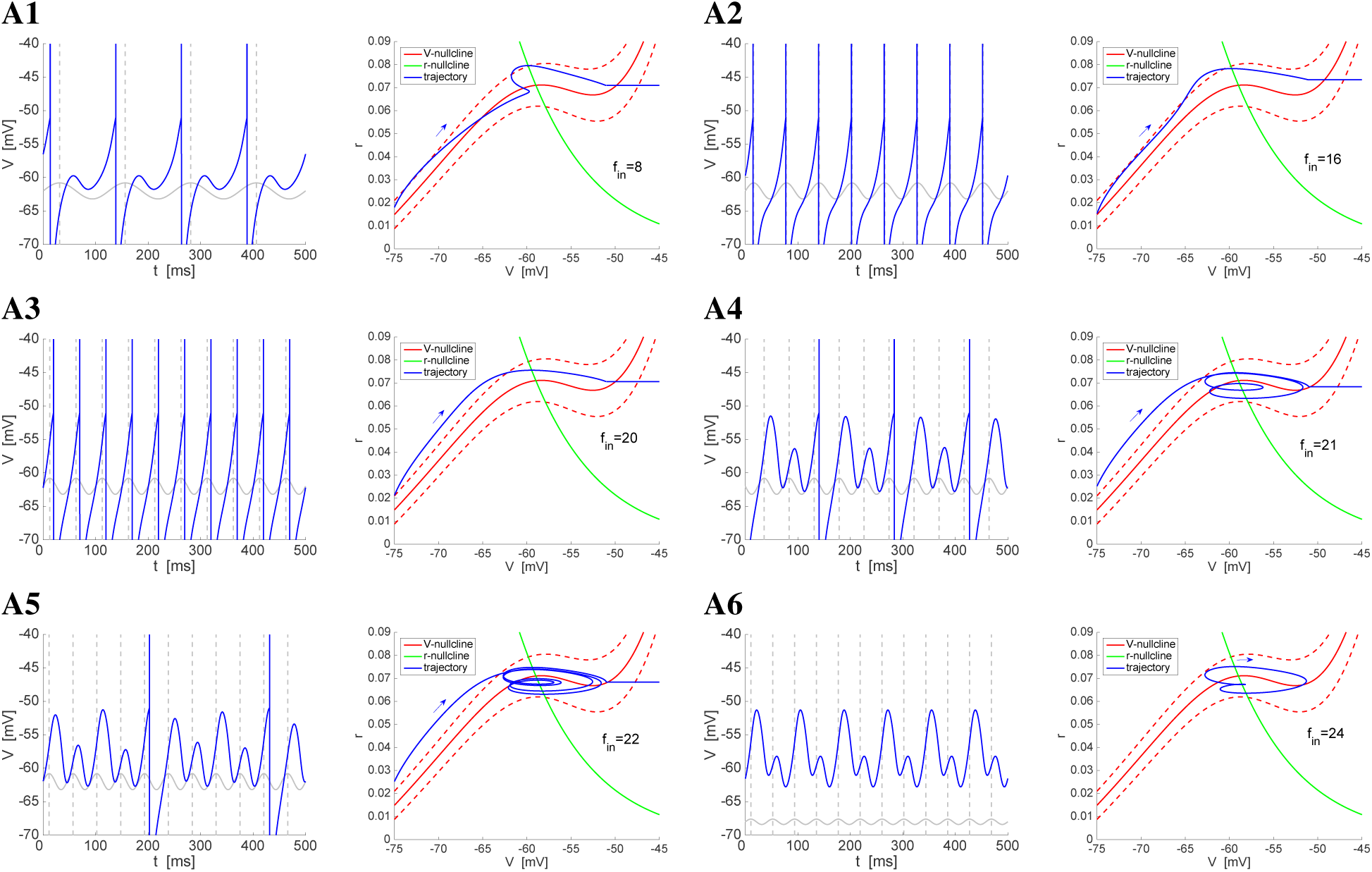
Suprathreshold response of the *I_h_*+*I_Nap_* model 2 to sinusoidal inputs for representative parameter values. *Voltage traces (left panels) and phase-plane diagrams (right panels) for A_in_* = 0.45 *and representative values of f_in_. Parameter values are as in Figs. 10 and 11 (and Figs. 7-B and Fig. 8-C) with two exceptions: The r-nullcline was shifted to the right to accommodate the change in I_app_ that shifted the V-nullcline up*. **Left panels:** *The solid-gray curves are caricatures of the sinusoidal inputs. The dashed-gray vertical lines at the peaks of the sinusoidal inputs indicate* Φ*_spk_* = 0 *(zero phase-shift)*. **Right panels:** *The dashed-red curves are the V-nullclines displace* ±*A_in_ units above and below the V-nullcline for the autonomous system (solid-red). They indicate the boundaries of the cyclic displacement of the V-nullcline as time progress due to the sinusoidal input. The V-nullcline reaches its lowest and highest levels (dashed-red curves) at a quarter and three quarters of each cycle respectively.The arrow indicates the direction of motion of the trajectory from its reset point to the spiking regime. We used the following parameter values: G_L_* = 0.3, *G_p_* = 0.09, *G_h_* = 1.5, *I_app_* = −1.2, *D* = 0, *V_th_* = −51, *V_rst_* = −75, *r_rst_* = 0.

Our results, presented in Figs. 14 to 17, show that our findings discussed in the previous sections persists for the changes in (*V_rst_, r_rst_*). There are some differences between the patterns obtained for the two different sets of reset values for the two models, but these differences are rather quantitative than qualitative. For example, model 1 has almost no 3:1 response patterns (Fig. 14, right panels). However, the 2:1 response patterns are generate by similar mechanisms as the ones described above (Fig. 16). In addition, model 2 displays response bursting patterns for high enough values of *A_in_* and intermediate input frequencies (Fig. 15-D and -E, black dots). This bursts are generated on top of 1:1 subthreshold oscillations (Fig. 17-B2 and -B3).

**Figure 14:**
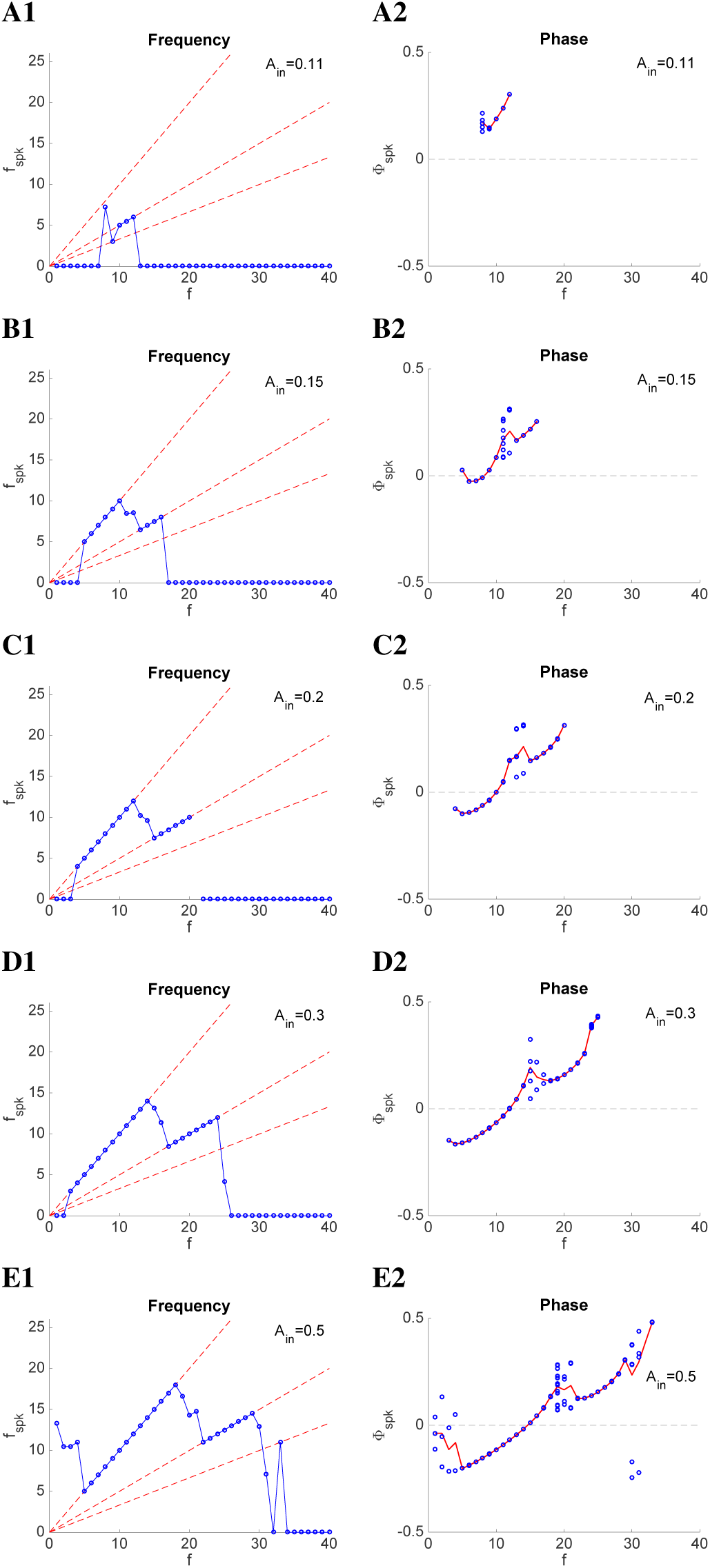
Suprathreshold response of the *I_h_*+*I_Nap_* model 1 to sinusoidal inputs for representative parameter values *(as in Figs. 7-A)*. **Left panels:** *Spike-frequency diagrams. The output spike frequency *f_spk_* is the inverse of the average length of the interspike intervals. The dashed-red lines (from top to bottom) indicate the 1:1, 2:1, and 3:1 output spikes versus input cycle patterns, respectively*. **Right panels:** *Spike-phase diagrams. The output spike phase* Φ*_spk_ (blue dots) was computed as the difference between the output spike-time and the closest input peaktime normalized by the cycle length*.Φ*_spk_* = 0 *for spikes at the input peak and* Φ*_spk_* = ±0.5 *for spikes at the immediate input troughs. The red line indicates the average* Φ*_spk_ for each input frequency. We used the following parameter values: G_L_* = 0.5, *G_p_* = 0.5, *G_h_* = 1.5, *I_app_* = −2.5, *D* = 0, *V_th_* = −45, *V_rst_* = −52, *r_rst_* = 0.05.

**Figure 15:**
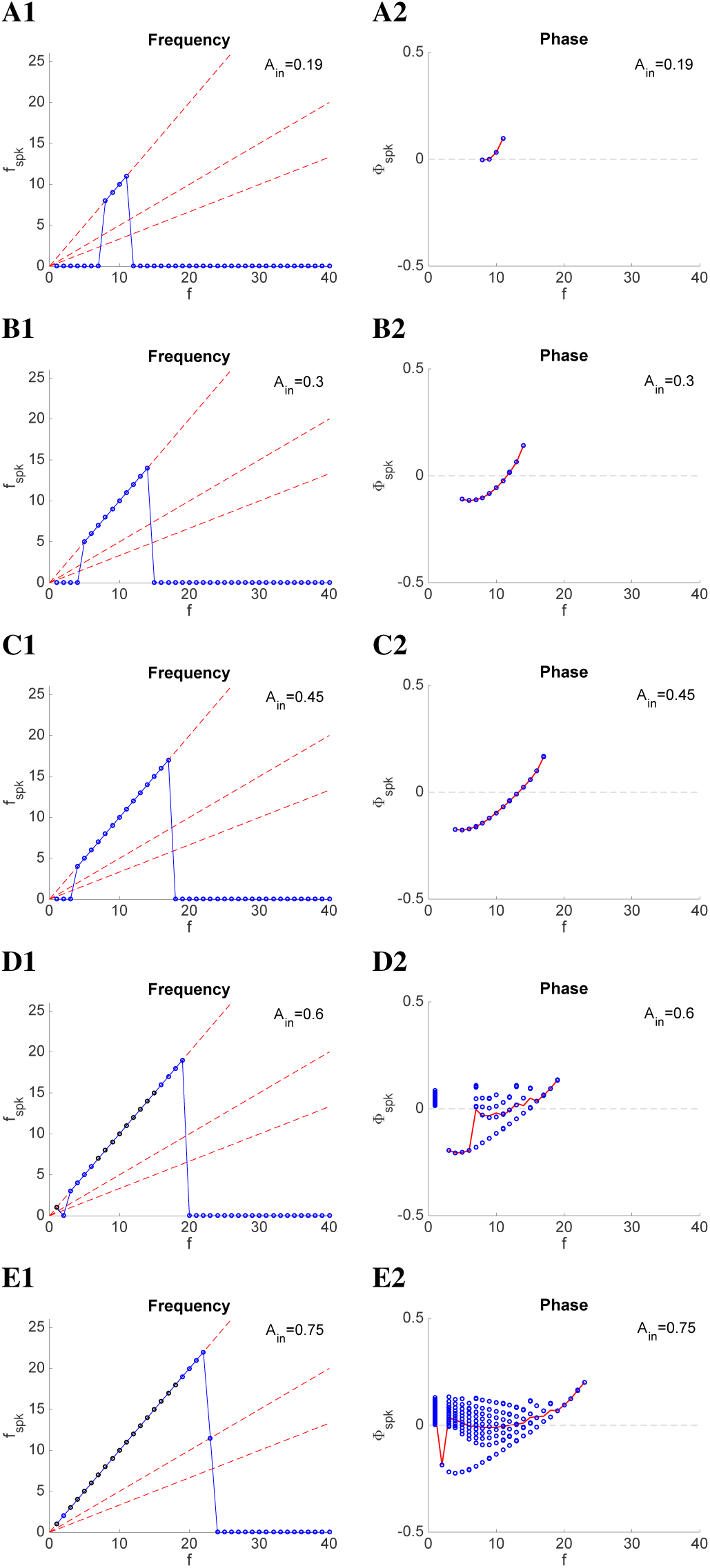
Suprathreshold response of the *I_h_*+*I_Nap_* model 2 to sinusoidal inputs for representative parameter values *(as in Figs. 7-B)*. **Left panels:** *Spike-frequency diagrams. The output spike frequency *f_spk_* is the inverse of the average length of the interspike intervals. The dashed-red lines (from top to bottom) indicate the 1:1, 2:1, and 3:1 output spikes versus input cycle patterns, respectively. The black dots indicates bursting behavior at a much higher intra-burst frequency*. **Right panels:** *Spike-phase diagrams. The output spike phase* Φ*_spk_ (blue dots) was computed as the difference between the output spiketime and the closest input peak-time normalized by the cycle length*. Φ*_spk_* = 0 *for spikes at the input peak and* Φ*_spk_* = ±0.5 *for spikes at the immediate input troughs. The red line indicates the average* Φ*_spk_ for each input frequency. We used the following parameter values: G_L_* = 0.3, *G_p_* = 0.08, *G_h_* = 1.5, *I_app_* = 0.3, *D* = 0, *V_th_* = −51, *V_rst_* = −52, *r_rst_* = 0.035.

**Figure 16:**
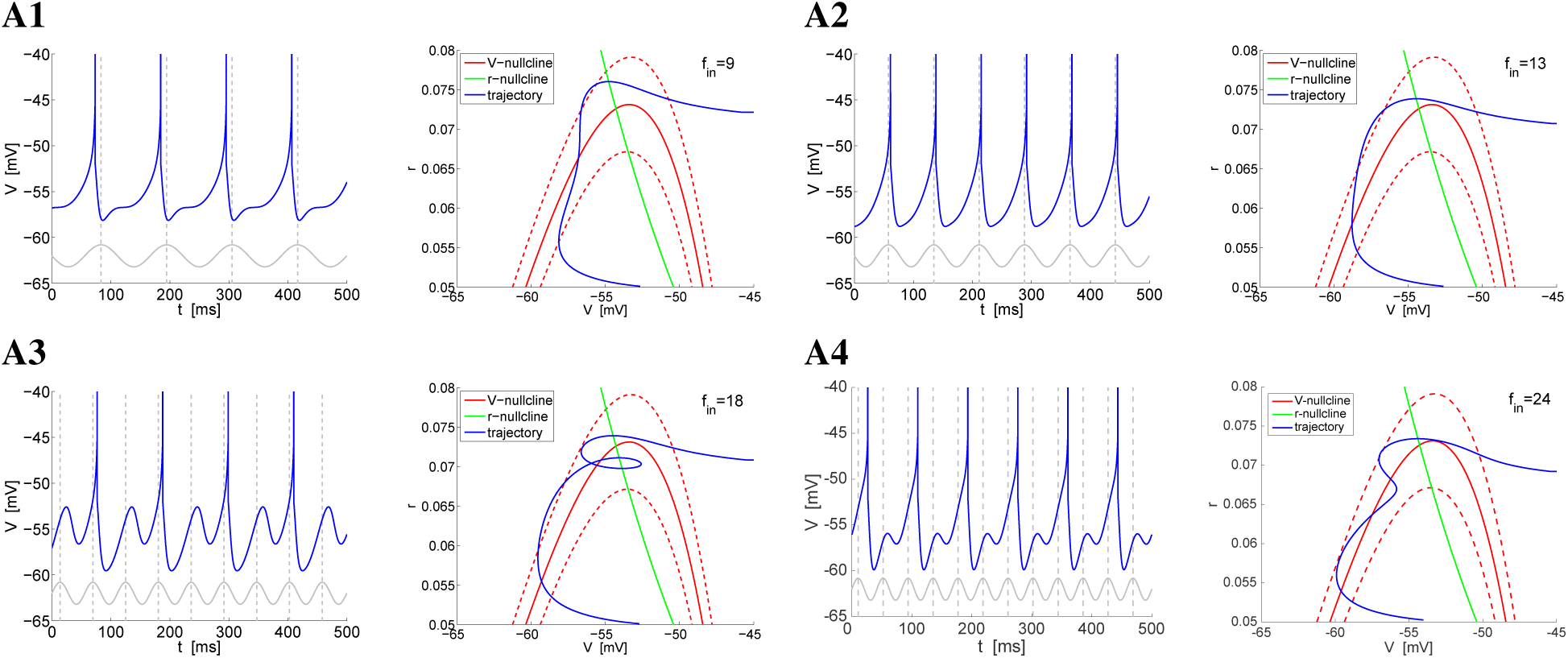
Suprathreshold response of the *I_h_*+*I_Nap_* model 1 to sinusoidal inputs for representative parameter values and *A_in_* = 0.3. *Parameter values are as in Figs. 7-A and Fig. 8-D*. **A.** *Voltage traces (left panels) and phase-plane diagrams (right panels) for representative values of the input frequency f_in_*. **Left panels:** *The solid-gray curves are caricatures of the sinusoidal inputs. The dashed-gray vertical lines at the peaks of the sinusoidal inputs indicate* Φ*_spk_* = 0 *(zero phase-shift)*. **Right panels:** *The dashed-red curves are the V-nullclines displace* ±*A_in_ units above and below the V-nullcline for the autonomous system (solid-red). They indicate the boundaries of the cyclic displacement of the V-nullcline as time progress due to the sinusoidal input. The V-nullcline reaches its lowest and highest levels (dashed-red curves) at a quarter and three quarters of each cycle respectively. The arrow indicates the direction of motion of the trajectory from its reset point to the spiking regime. We used the following parameter values: G_L_* = 0.5, *G_p_* = 0.5, *G_h_* = 1.5, *I_app_* = −2.5, *D* = 0, *V_th_* = −45, *V_rst_* = −52, *r_rst_* = 0.05.

**Figure 17:**
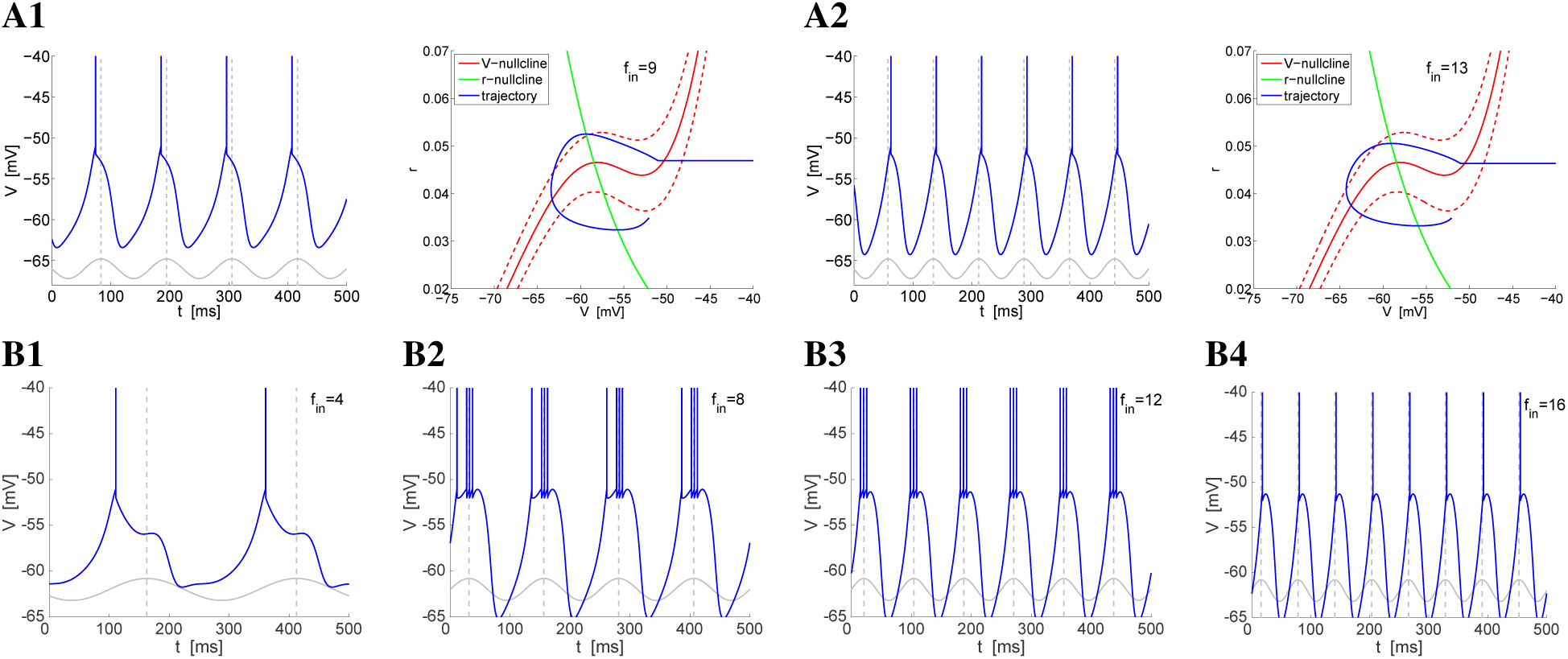
Suprathreshold response of the *I_h_*+*I_Nap_* model 2 to sinusoidal inputs for representative parameter values *(as in Figs. 7-B and Fig. 8-C)*. **A.** *A_in_* = 0.45. *Voltage traces (left panels) and phase-plane diagrams (right panels) for representative values of the input frequency f_in_*. **Left panels:** *The solid-gray curves are caricatures of the sinusoidal inputs. The dashed-gray vertical lines at the peaks of the sinusoidal inputs indicate* Φ*_spk_* = 0 *(zero phase-shift)*. **Right panels:** *The dashed-red curves are the V-nullclines displace* ±*A_in_ units above and below the V-nullcline for the autonomous system (solid-red). They indicate the boundaries of the cyclic displacement of the V-nullcline as time progress due to the sinusoidal input. The V-nullcline reaches its lowest and highest levels (dashed-red curves) at a quarter and three quarters of each cycle respectively.The arrow indicates the direction of motion of the trajectory from its reset point to the spiking regime*. **B.** *A_in_* = 0.6. *Voltage traces. We used the following parameter values: G_L_* = 0.3, *G_p_* = 0.08, *G_h_* = 1.5, *I_app_* = 0.3, *D* = 0, *V_th_* = −51, *V_rst_* = −52, *r_rst_* = 0.035.

## 4 Discussion

Neuronal models have been classified using different criteria. The most natural one is based on the identify of the participating ionic currents [58, 67]. According to this biophysical classification the *I_h_* + *I_Nap_* and *I_Ks_* + *I_Nap_* models are different. An alternative classification scheme is based on the geometric and dynamic properties of the phase-space diagrams. In the subthreshold voltage regime, the voltage nullclines (or nullsurfaces) are typically quasilinear, parabolic-like or cubic-like [58, 62, 64, 68]. According to this classification, the parabolic- and cubic-like *I_h_* + *I_Nap_* models are different and so they are the corresponding *I_Ks_* + *I_Nap_* models, and the parabolic-like *I_h_* + *I_Nap_* and *I_Ks_* + *I_Nap_* belong to the same class and so they do the corresponding cubic-like models [58]. This classification is useful to understand the similarities and differences among the various mechanisms underlying the generation of STOs and other patterns that are, at least, partially controlled by the neuron’s subthreshold currents. In previous work, we compared the STO properties of these models and we showed that while some of these properties depend on the specific types of ionic currents involved, and they are different for the *I_Nap_* + *I_h_* and the *I_Nap_* +*I_Ks_* models, others depend on the type of voltage nullclines involved and are shared by models with different ionic currents. This suggested that the responses to oscillatory inputs of parabolic- and cubic-like models having the same type of ionic currents might be different.

We set out to examine these issues in the context of the parabolic- and cubic-like *I_h_* + *I_Nap_* models investigated in [58] (referred to as model 1 and 2 respectively in this paper). The salient outcomes of our study are (i) the identification of the similarities and differences in the subthreshold and spiking resonant properties between the parabolic- and cubic-like model versions, (ii) the explanation of the mechanisms that underlie these phenomena, and (iii) the identification of conditions under which the subthreshold resonant properties are communicated to the spiking regime. Overall, our results show that the effective time scales that operate in the subthreshold regime to generate intrinsic STOs, MMOs and subthreshold resonance do not necessarily determine the spiking response to oscillatory inputs to be in the same frequency band.

Both models exhibit subthreshold resonance in the theta frequency band, but for values of the input amplitude *A_in_* close to the threshold for spike generation the voltage response is nonlinearly amplified (the impedance increases with increasing values of *A_in_*) for the parabolic-like model as predicted in [61], while it is nonlinearly attenuated (the impedance decreases with increasing values of *A_in_*) for the cubic-like model.

For low enough values of *A_in_* the two models exhibit both evoked and output spiking resonance in the theta frequency band, implying that the subthreshold resonances in both models are communicated to the suprathreshold voltage regime. However, for higher values of *A_in_* the evoked spiking resonance disappears in both models reflecting the fact that the broader input frequency bands cause the voltage responses to be above threshold. However, the parabolic model still exhibits output spiking resonance for these values of *A_in_*, while the cubic-like model shows 1:1 entrainment where each input cycle generates either a single spike or a burst (in the latter case it would be technically more appropriate to speak of N:1 entrainment with N ≥ 1). The output spiking resonance in the parabolic-like model involves the generation of MMOs by a cycle skipping mechanism. This cycle skipping mechanism is absent in the cubic-like model for the relevant input frequencies. However, based on our results we cannot rule out its presence for the higher input frequencies in other parameter regimes. More research is needed to determine whether the transition from 1:1 entrainment to no spiking is abrupt or gradual (involving MMOs) through a small input frequency range.

The parabolic- and cubic-like *I_h_* + *I_Nap_* models we investigate in this paper have been used to study the subthreshold properties of medial entorhinal cortex layer II stellate cells [61, 62, 64], are similar to other models used with the same purpose [7], and are representative of a more general class of models involving the interplay of these two currents. Moreover, they are representative of a more general class of models involving the interplay of two ionic currents having a fast amplifying and a slow resonant gating variables (e.g., *I_Ks_* + *I_Nap_* models). Therefore, our findings have implications for a generic class of systems exhibiting subthreshold and spiking resonance, such as the *I_Ks_* + *I_Nap_* models mentioned above, exhibiting resonances in a broad range of frequency bands. Importantly, models having multiplicative amplifying and resonant gating variables (e.g, L-type high-threshold calcium currents) are excluded from this group since their voltage nullclines are typically cubic-like and rarely (or never) parabolic-like. The predictions generated by our results can be tested experimentally in a variety of systems. These experiments may also serve to discriminate between different types of nonlinearities present in the biophysical models.

Most of the results presented in this paper are based on the time constant for medial entorhinal cortex stellate cells (*τ_r_* = 80) used in [66] for the fast component of *I_h_*, and are therefore restricted to the theta frequency band. However, *τ_r_* for *I_h_* is highly variable across cells and species. Changes in the values of time constants in neuronal models not only affect the oscillation frequency, but also other properties such as the oscillation amplitude. In fact, decreasing values of *τ_r_* causes the cells’ to behave closer to their linearization [61]. We examined the effects of changes in *τ_r_* on the main results presented in this paper by using two representative values above and below *τ_r_* = 80: *τ_r_* = 200 and *τ_r_* = 40 (close to the value reported for CA1 pyramidal cells).

The differences between the two type of models persist for a large range of values of *τ_r_*, but there are some quantitative differences both between the two models and among the different values of *τ_r_* for each model. In all cases, for low enough suprathreshold values of *A_in_* both models exhibit evoked and output spiking resonance in frequency bands that coincide with the subthreshold resonant frequency band for each value of *τ_r_*. For the parabolic-like model and *τ_r_* = 200 the output frequency band remains bounded, roughly coincides with the subthreshold frequency band for larger values of *A_in_*, and is narrower than the output frequency band for *τ_r_* = 80. For *τ_r_* = 40 the cycle skipping mechanism giving raise to the MMO response patterns is not strong enough to prevent the (1:1) entrainment for large enough values of *A_in_*. The output frequency band in these cases is therefore larger than the subthreshold resonant frequency band, and therefore output resonance is no longer observed for largest values of *A_in_* we used. For the cubic-like model and *τ_r_* = 200 a cycle skipping mechanism giving rise to MMO response patterns causes the (1:1) entrainment to be weaker than for *τ_r_* = 80, but still the entrainment is strong enough to produce a relatively large output frequency band, well above the subthreshold frequency band. Evoked resonant is present for a larger range of values of *A_in_* as compared to *τ_r_* = 80, but it disappears for values of *A_in_* beyond this range. For *τ_r_* = 40, the (1:1) entrainment is as for *τ_r_* = 80 and there is no output spiking resonance, except for the low enough subthreshold values of *A_in_* for which, as mentioned above, there are both evoked and output spiking resonance. In these cases, the entrainment involves MMO patterns (2:1 entrainment). This highlights the important role played by the time constant of the resonant current in determining the resonant spike response patterns. Future research should address this issue.

The differences between the parabolic- and cubic-like models is not restricted to the shapes of the *V*-nullclines but include also the spiking mechanisms. The parabolic-like model describes the onset of spikes in addition to the subthreshold dynamics (Fig. 5-A3). Spikes occur when the trajectory moves along a fast direction to the right of the *V*-nullcline and escapes the subthreshold regime. The voltage threshold only indicates the occurrence of spikes and is not part of the spiking mechanism. In contrast, the voltage threshold is part of the spiking mechanism in the cubic-like models (Fig. 5-B5). A more natural spiking mechanism for the cubic-like model would involve an additional ionic current, therefore making it three-dimensional. The addition of the standard spiking currents (transient sodium and delayed rectifier potassium) to the parabolic-like *I_h_* + *I_Nap_* model that would generate “natural” spikes is not expected to produce qualitative changes in our results, but perhaps minor quantitative differences in the resonant frequency bands and other magnitudes. Whether or not this is the case for the cubic-like *I_h_* + *I_Nap_* model is an open question.

The results discussed in this paper highlight both the complexity of the suprathreshold responses to oscillatory inputs in neurons having resonant and amplifying currents with different time scales and the fact that the identity of the participating ionic currents is not enough to predict the resulting patterns, but additional dynamic information captured by the geometric properties of the phase-space diagram is needed. This has implications for mechanistic studies on suprathreshold (firing rate and spiking) resonances [2, 6, 78, 79] as well as other types of preferred frequency responses to oscillatory inputs including phase-locking [47, 80, 81], synchronization [82, 83], synaptic [18, 84–86], and network [51–53, 87–95] ones.

## Acknowledgments

This work was partially supported by the NSF grants DMS-1313861 and DMS-1608077 (HGR). The author thanks Eran Stark for useful comments.

